# Histone succinylation directly inhibits Jumonji domain demethylases and stabilizes repressive chromatin states

**DOI:** 10.64898/2026.05.29.728167

**Authors:** Sarah Graff, Stephanie Stransky, Isaac Kraz, Kheerthana Duraivelan, Alexander Poppel, Adam Chatoff, Nathaniel W. Snyder, Scott Garforth, David Shechter, Juan Pablo Maianti, Simone Sidoli

## Abstract

Herein we uncover a relationship between histone succinylation and Jumonji (JmjC) domain-containing histone demethylases. We used quantitative proteomics and peptide pull-down assays to identify JmjC demethylases as candidate interactors with succinylated histone peptides. Succinyl-lysine peptides bind and inhibit the catalytic activity of JmjC demethylases in a dose-dependent manner. This includes KDM4D and KDM6B, which are responsible for the removal of the silencing marks H3K9me2/3 and H3K27me2/3. Supraphysiological sodium succinate treatment of HepG2/C3A cells increased the relative abundance of histone succinylation, H3K9me2/3, and H3K27me2/3. CUT&Tag and ChIP–mass spectrometry revealed the co-occurrence of succinylation with these repressive methylation marks, in addition to reduced transcriptional output. This work establishes a novel mechanistic link between metabolite abundance and chromatin regulation and suggests a role for histone succinylation in the maintenance of heterochromatin.

## INTRODUCTION

Histone post-translational modifications (PTMs) play a crucial role in mediating chromatin dynamics, which lead to downstream effects on gene expression without alterations to the underlying DNA sequence^1^. The sources of these PTMs are chemical adducts derived from cellular metabolites, and the abundances of these metabolites frequently correlate with the relative abundance of the respective histone PTMs^2^. Therefore, histone PTMs reflect a link between cellular metabolism and gene expression. The most frequently observed histone PTMs have been the subject of significant research, such as lysine acetylation (Kac) and methylation (Kme), for which the binding partners and functional consequences on chromatin state are well understood. Histone PTMs can function in concert with each other in a phenomenon known as “crosstalk”, where specific combinations of modifications enhance or restrict binding of chromatin modifiers and transcription factors^3^. In addition to these well-characterized and abundant histone PTMs, there are also many other low-abundance histone acyl modifications whose functional role and link to cellular metabolism has yet to be fully described^4^.

Histone lysine succinylation (Ksu), first discovered in 2011 in multiple eukaryotic species, is one of these poorly characterized PTMs^5^. Succinylation of lysine adds a mass shift of +100 Da and alters the side-chain charge state from +1 to -1 at physiological pH^6^. In comparison, lysine acetylation is a smaller adduct (+42 Da) and neutralizes the positive charge of the lysine side-chain. This weakens the histone electrostatic interaction with negatively charged DNA, promoting open chromatin and facilitating active transcription^7^. Since lysine succinylation is approximately twice the size of acetylation, and negatively charged, it has been theorized to have a more significant effect on the formation of open chromatin. Chromatin profiling experiments have shown that histone succinylation is enriched at gene promoters and transcription start-site regions^8^. Moreover, histone succinylation correlates with histone PTMs associated with active gene transcription, such as H3K4me3, as well as increased transcription of genes when both PTMs are present ^9,10^. Despite the evidence for synergy between succinylation and markers of euchromatin, succinylation lacks epigenetic readers in common with histone acetylation. Specifically, bromodomains, which bind Kac within a hydrophobic pocket, are routinely found to be unable to bind Ksu, suggesting a distinct functional role for this PTM^11^. Several histone acetyltransferases have also been shown to act as succinyltransferases, including p300, KAT2A, and HAT1^8,12,13^. The de-acetylase sirtuin proteins 5 and 7, as well as the HDACs 1/2/3, have been shown to have desuccinylase activity^10,14,15^.

The acyl donor for succinylation is succinyl-coenzyme A (CoA), a central metabolite in the Kreb’s/tricarboxylic acid (TCA) cycle.^16^ Succinyl-CoA can be generated by activation or exchange of succinate with CoA via diverse enzymes including succinyl-CoA synthetase, as well as from α-ketoglutarate (α-KG) and CoA via the α-KG dehydrogenase (OGDH) complex^17,18^. Recent reports have indicated non-mitochondrial localization of both succinyl-CoA synthetase and OGDH^12,19,20^. Thus, succinyl-CoA formation relies on α-KG, as well as succinate abundance. Due to its carboxylic acid in proximity of the thioester, succinyl-CoA can undergo self-hydrolysis to form a reactive anhydride intermediate. Succinic anhydride is a non-enzymatic modifier of lysine residues, correlating the abundance of protein succinylation and the pool of available succinyl-CoA^21^. Non-enzymatic modification appears to be the major source of histone succinylation, given that succinyl-CoA is a structurally unsuitable co-factor for the catalytic site of histone acetyltransferase enzymes. The stochastic process of non-enzymatic histone succinylation accumulates in sites closer to the C-terminus, or core, of the histone proteins, while acyltransferases target the N-terminal tails of histones^22^.

A known nuclear source for succinate or the precursor α-KG is the activity of a group of chromatin modifiers, the JmjC domain-containing iron- and α-KG-dependent demethylase family. JmjC domain demethylases target four lysine residues on the histone H3 tail, oxidatively removing both silencing and activating marks. Each demethylation reaction requires the decarboxylation of α-KG as a cofactor, which is converted to succinate^23^. The relationship between metabolite abundance and chromatin state also plays a role in JmjC demethylase activity, through inhibition by succinate, as well as similar TCA metabolites like fumarate. Under elevated succinate conditions, this leads to an increase in di- and tri-methylation on the H3 tail^24^.

In this study, we identify a previously uncharacterized interaction between histone succinylation and JmjC domain demethylases. Using proteomic and biochemical approaches, we show that Ksu histone peptides inhibit the demethylase activity of these enzymes, resulting in increased methylation at canonical lysine target sites under conditions of elevated histone succinylation. Furthermore, we provide evidence that elevated histone succinylation induced through long-term succinate treatment is associated with a global reduction in transcriptional output, consistent with the accumulation and spread of heterochromatic lysine methylation.

## RESULTS

### Histone succinylation is a quantifiable and metabolically responsive dynamic post-translational modification

Previous work has shown that the availability of metabolite donors for acyl modification correlates with the abundance of acyl-lysine PTMs, both *in chemico* and *in vitro*^25^. Based on this rationale, we designed an experimental scheme to modulate histone succinylation. We treated HepG2/C3A cells with 10 mM sodium succinate and quantified changes in the abundance of histone lysine succinylation using nano liquid chromatography coupled online with mass spectrometry (LC-MS) followed by signal extraction using the software EpiProfile 2.0^26^. We used both flat cell culture (2D) and spheroids produced by 3D cell culture, which replicate more slowly and are more metabolically similar to liver physiology^27^. We found that succinate treatment for 48 hours in 2D culture, and one week in 3D cell culture, both significantly increase the global levels of histone succinylation in HepG2/C3A cells (**Fig. 1A-B**). We hypothesize that the slowed growth rate in the 3D cell culture may allow for unique PTM distributions as compared to 2D cell culture, by giving time for PTMs with slower turnover to accumulate on histones (**Fig. 1C**)^28,29^. We found that, irrespective of succinate treatment, the residue most frequently succinylated in 2D cell culture was H3K64 (**Fig. 1C**). Notably, H3K64 is located within the core of the histone protein, but still solvent accessible (Rasmol model, **Fig. 1D**). In contrast, the most frequently succinylated residues in succinate-treated spheroids were in the core of histone H4 (K77 and K79). These results are consistent with previous observations of non-enzymatic histone succinylation occurring primarily on the histone core^22^. Non-enzymatic histone succinylation is hypothesized to be driven by elevated levels of succinyl-CoA in the nucleus^30^, which we measured by using fractionation and metabolomics^30^. We found that, following treatment with succinate, HepG2/C3A cells displayed an increase of succinyl-CoA in the nucleus but not in the non-nuclear fraction (**Fig. 1E**). This data supports the model that non-enzymatic succinylation is the primary mechanism behind the observed increase in global histone succinylation^22^. The 48-hour timepoint of succinate treatment lacked any conspicuous cellular phenotype (**Fig. 1F**). However, continuous treatment of flat cell culture with succinate (up to 144h) slowed cell growth rate, with unchanged viability (**Fig. 1F, Fig. S1B**).

**Figure 1.**
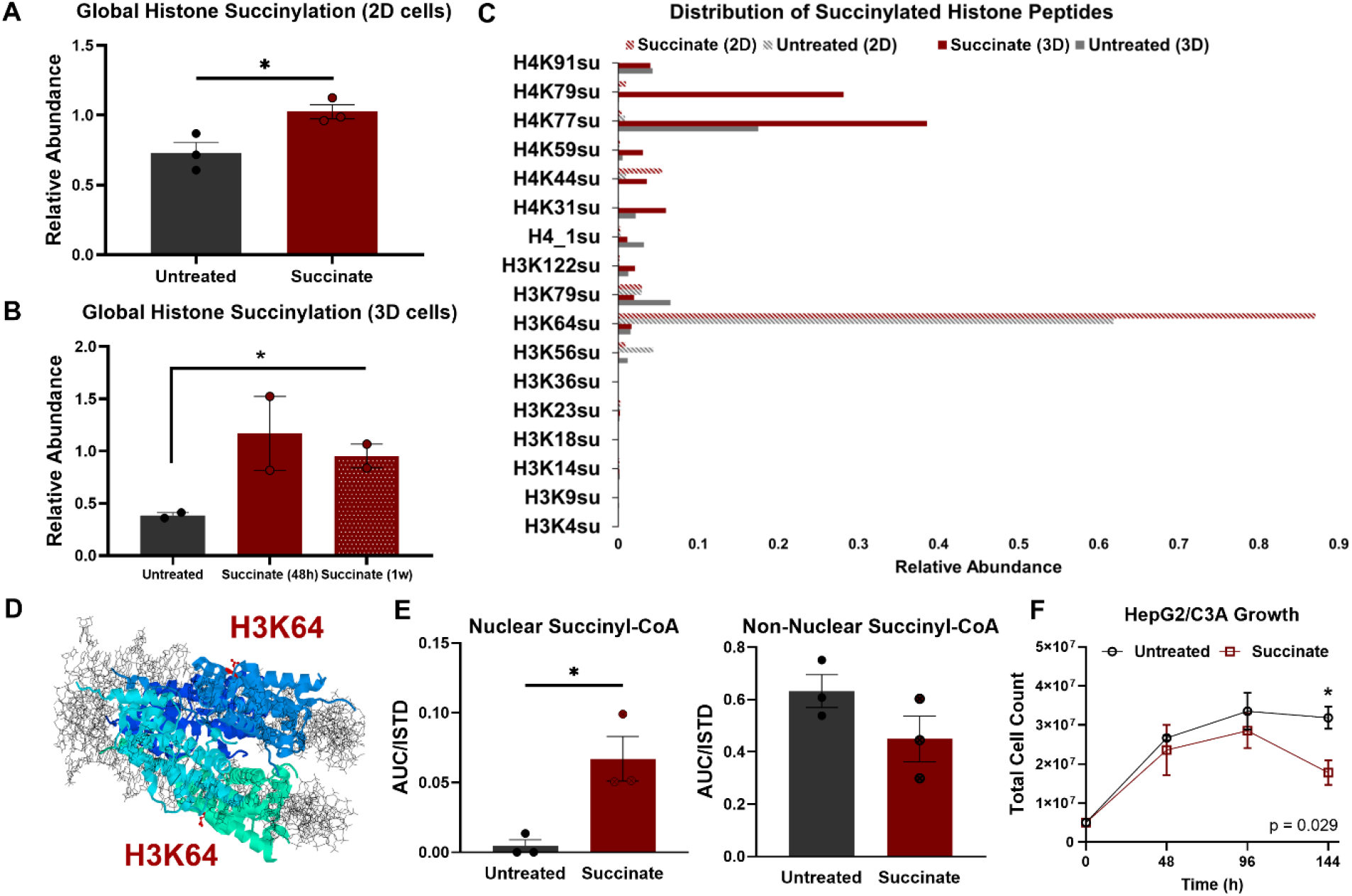
Histone succinylation is a dynamic histone post-translational modification. (A) Global histone succinylation measured by LC-MS/MS in HepG2/C3A cells following 48 hours of sodium succinate treatment. (B) Global histone succinylation in 3D cultured HepG2/C3A spheroids following 48 hours and 1 week of sodium succinate treatment. (C) Comparison of the relative abundance of succinyl-lysine residues in H3 and H4, in either treated or untreated, cultured cells (2D) or spheroids (3D). (D) Structure of the nucleosome with H3K64 residues highlighted to demonstrate solvent accessibility. (E) Abundance of succinyl-CoA in the nuclear fraction (left) and non-nuclear fraction (right) following sodium succinate treatment. (F) Total count of HepG2/C3A cells treated with 10 mM sodium succinate continuously for 48 to 144 hours, cell count taken every 48 hours. *Data are shown as mean ± SEM; * p-value <0*.*05 in two-tailed Student’s t-test*.

### Jumonji domain demethylases interact with histone lysine succinylation

We used a biotinylated peptide pull-down mass spectrometry assay to identify candidate protein interactors with histone lysine succinylation. Synthetic histone peptides displayed on streptavidin beads were incubated with the nuclear fraction of HepG2/C3A cell lysate. The nuclear proteins enriched by the modified peptides were analyzed with a standard bottom-up proteomic workflow (**Fig. 2A**). As expected, the acetylated histone peptide (positive control) enriched a large number of known binder proteins (**Fig. 2B**). We compared the enrichment of bromodomain-containing proteins to the synthetic peptides and demonstrated that these proteins do not bind to Ksu peptides in this pull-down experiment, supporting previous findings^11^ (**Fig. 2C**). Notably, the bromodomain proteins displayed greater enrichment to the unmodified histone peptide than to the Ksu peptide (**Fig. 2D**). We then searched for protein families specifically enriched by the Ksu peptide and identified the JmjC domain histone demethylases (**Fig. 2E-F**). JmjC demethylases are Fe(II)/αKG-dependent dioxygenases in which αKG coordinates the catalytic iron within an evolutionarily conserved pocket required for oxidative demethylation (**Fig. 2G**). Previous studies have shown that the enzyme’s product, succinate, can occupy and inhibit the αKG co-factor site of Jumonji domain demethylases^31^. Daminozide, a pesticide with structural similarity to a succinyl moiety, has also been shown to inhibit the activity of KDMs^32^. The enrichment of evolutionarily distinct JmjC domain demethylases to succinylated peptides was further supported by AlphaFold and Boltz2 models (**Table S2–S3**), suggesting that Ksu could similarly occupy the αKG co-factor site in a manner that appears broadly allowed by JmjC domains (**Fig. 2G–H, Fig. S2–S3**). Consistent with this rationale, proteomic analysis of the chromatin fraction of succinate-treated HepG2/C3A cells revealed significant enrichment of multiple JmjC domain demethylases compared to untreated cells (**Fig. 2I**). These novel observations suggest that elevated histone succinylation leads to previously unknown chromatin-binding interactions with JmjC domain demethylases in ways that could directly or indirectly affect gene expression.

**Figure 2.**
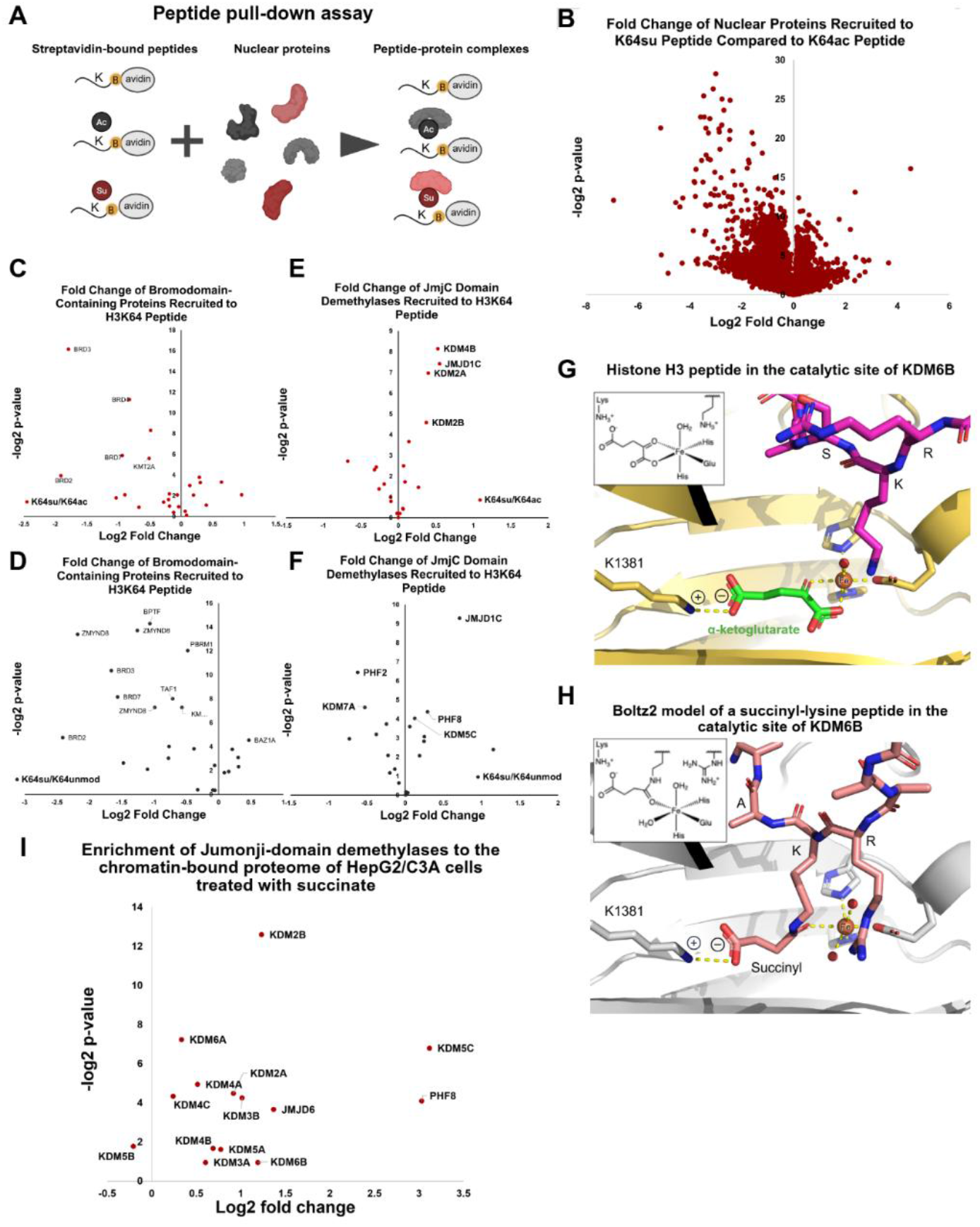
Jumonji domain lysine demethylases exhibit an affinity for succinyl-lysine histone peptides. (A) Schematic of peptide pull-down technique used to identify potential protein interactors with histone succinylation. (B) Volcano plot of fold change and significance of enrichment of nuclear proteins from HepG2/C3A cells to a Ksu synthetic peptide vs. Kac synthetic peptide. (C) Volcano plot of fold change and significance of enrichment of bromodomain proteins to Ksu synthetic histone peptide as compared to the Kac peptide, and (D) Ksu synthetic histone peptide as compared to an unmodified peptide. (E) Volcano plot of fold change and significance of enrichment of JmjC domain demethylases to Ksu synthetic histone peptide as compared to an Kac peptide and (F) Ksu synthetic histone peptide as compared to an unmodified peptide. (G) Crystal structure (PDB 6F6D) of an unmodified histone H3 peptide in the catalytic site of KDM6B. (H) Boltz2 model of a Ksu peptide (AARK(succinyl)A) in the catalytic site of KDM6B. (I) Enrichment of JmjC-domain demethylases to the chromatin-bound proteome of HepG2/C3A cells treated with 10 mM sodium succinate for 48 hours. *A -log2 p-value greater than 4 indicates significant enrichment*.

### Succinyl-lysine peptides inhibit the demethylation activity of Jumonji domain demethylases

To characterize the interaction between JmjC demethylases and lysine PTMs, we synthesized acylated lysine peptides (**Fig. 3A**) and tested these in a panel of commercially available assays at Reaction Biology Corporation and Eurofins. These assays use the principles of AlphaLISA (amplified luminescent proximity homogeneous) immunoassay, or homogeneous time-resolved Förster resonance energy transfer (HTRF/TR-FRET), to detect the loss of methyl groups on a biotinylated Kme3 histone substrate peptide (**Fig. 3B**). We chose a minimal 6-mer backbone for the synthetic peptides; A-A-R-K[modification]-S-A and specifically avoided hydrophobic residues to provide information on the structural basis of the PTM interaction with the enzyme. These assays identified two members of the JmjC family, KDM6B and KDM4D, that displayed dose-response inhibition of demethylase activity by Ksu synthetic peptides (**Fig. 3C-D**). In contrast, Kac synthetic peptides were unable to inhibit the catalytic activity of any assayed demethylases. It should be underscored that the succinyl-lysine PTM appears to be a generally modest driver of binding affinity, suggesting that the surrounding sequence and additional protein–protein interactions play a role in a biological context. We also tested eight additional JmjC domain demethylase assays and found similarly weak or undetectable inhibition with these peptides (**Fig. S4A-B)**. We also tested the ability of analogous lysine PTMs to inhibit the activity of the KDM6B and KDM4D JmjC domain demethylases (**Fig. 3A**). These PTMs are typically not detectable as histone modifications in human cells but may have biological relevance in other organisms^5,33–35^. Fumaryl- and maleyl-lysine represent conformationally locked *trans* and *cis* 4-carbon analogs of Ksu, respectively (**Fig. 3A**), and we also tested malonyl- and glutaryl-lysine as 3- and 5-carbon analogs, respectively. Fumaryl-lysine produced the strongest inhibition in KDM6B and KDM4D assays, corroborating our models of the PTM preferentially binding in the same extended conformation as αKG (**Fig. 2H**). However, all negatively charged acyl-peptides, including maleyl-lysine, routinely inhibited the assays to a greater extent than acetyl-lysine peptides (**Fig. 3E-F**). This is consistent with the spacious catalytic site of JmjC demethylases and the fact that our synthetic peptides lack hydrophobic anchoring residues for the substrate-binding site. Similarly, we observed that succinyl-lysine peptides were more potent than succinylated homo-lysine and ornithine peptides in KDM6B and KDM4D assays (**Fig. S4C**). LC-MS/MS analysis of the AlphaLISA reaction components post incubation with αKG and iron did not detect oxidation or hydrolysis of the succinyl-, fumaryl-, or maleyl-lysine peptides, suggesting these peptides are not competitive substrates (**Fig. S5A**). To allay any concerns that the succinyl-lysine peptides may interfere with AlphaLISA assay components, a synthetic sample of the biotinylated Kme2 products of KDM6B and KDM4D was spiked into the assays at Reaction Biology Corporation. The succinyl-lysine peptides did not inhibit these control assays (**Fig. S5B**).

**Figure 3.**
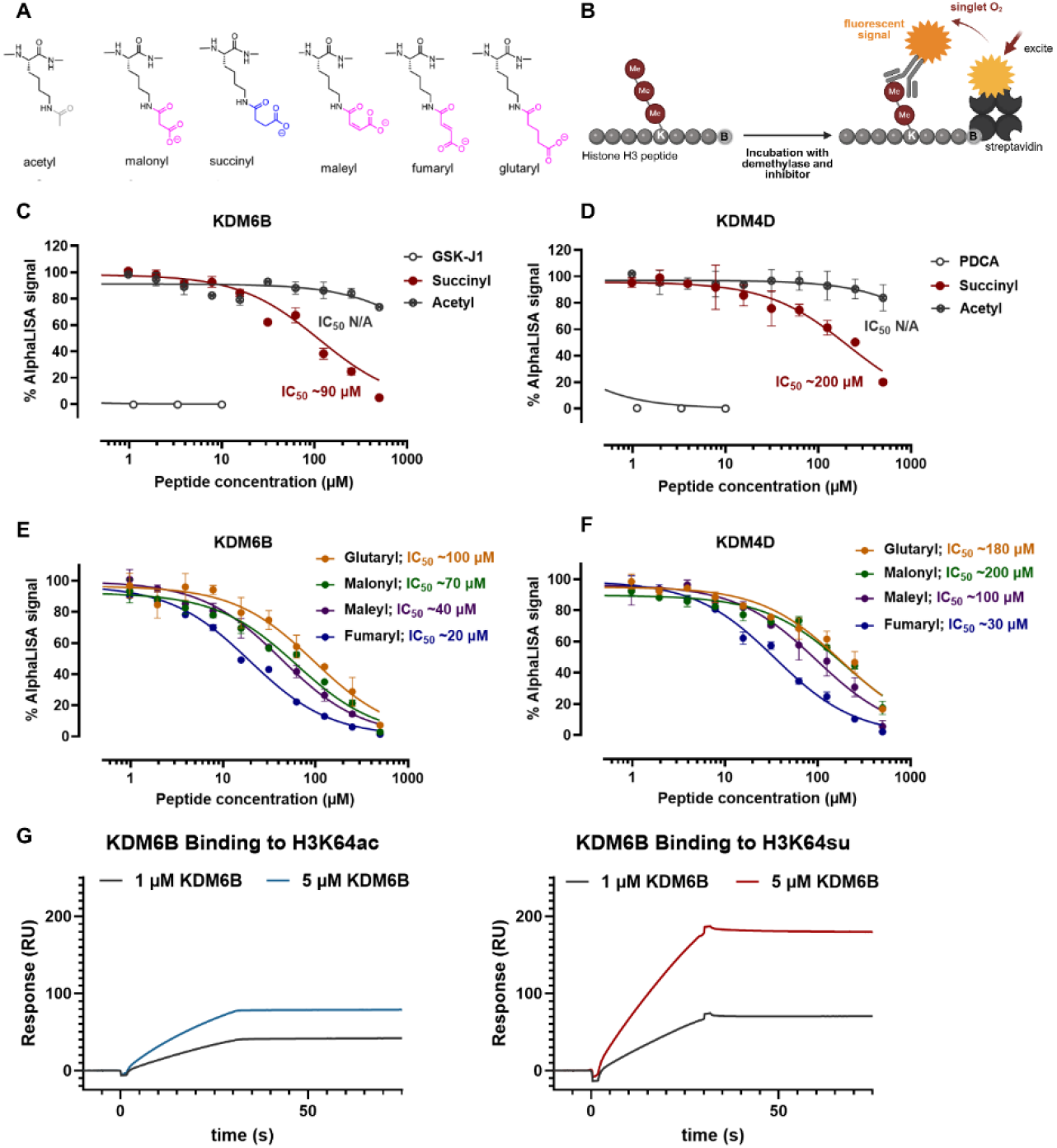
Succinyl-lysine synthetic histone peptides inhibit the demethylation reaction of Jumonji domain demethylases. (A) Structures of lysine PTMs tested in activity-based AlphaLISA assays. (B) Schematic of AlphaLISA demethylase activity assay procedure. JmjC domain demethylases are incubated with biotinylated tri-methylated histone H3 peptide substrates and proposed inhibitor, and demethylases activity is detected with an antibody specific to the demethylated product conjugated to an acceptor bead. Streptavidin donor beads bind to the biotinylated peptides and transfer a singlet oxygen to the antibody acceptor bead, producing a fluorescent signal. AlphaLISA assay of KDM6B (C) and KDM4D (D) activity in the presence of succinyl-lysine and acetyl-lysine histone-like peptide (GSK-J1 and PDCA are known KDM6B and KDM4D inhibitors). AlphaLISA assay of KDM6B (E) and KDM4D (F) activity in the presence of acylated lysine peptides. (G) Sensorgram of injections of KDM6B over acetyl-lysine (left) and succinyl-lysine (right) synthetic histone peptides. Data shown is an average of two replicate injections.

To further validate the interaction between histone succinylation and JmjC demethylases, we performed surface plasmon resonance (SPR) experiments using immobilized synthetic histone peptides. Biotinylated histone peptides modified with either succinyl-lysine (H3K64su) or acetyl-lysine (H3K64ac) were immobilized on the sensor surface, and KDM6B was injected as an analyte (**Table S1**). Under these conditions, we observed a robust dose-dependent binding response to the succinyl-lysine peptide consistent with specific binding (**Fig. 3G**). In contrast, the binding response to the acetyl-lysine peptide was lower and did not have the same dose-dependent response. These results indicate that Jumonji domain demethylases directly bind succinyl-lysine histone peptides, supporting a model in which lysine succinylation engages the enzyme cofactor- and substrate-binding pocket independently of catalysis.

### Increased global histone succinylation co-occurs with increased methylation at targets of Jumonji domain demethylases

The methyl-lysine substrates of human Jumonji-domain demethylases are well characterized and typically found on the histone H3 tail (e.g. K4, K9, K27, and K36 di-/tri-methylation) (**Fig. 4A**). These marks are typically associated with heterochromatin, except for H3K4me3, which correlates with active transcription. Previous studies examining the link between succinate and αKG-dependent enzymes found that elevated intracellular levels of succinate inhibited JmjC domain demethylases, leading to an increase in lysine di/tri-methylation at known target sites of these enzymes^16,24^. We treated HepG2/C3A cells with 10 mM sodium succinate to further examine the relationship between global histone succinylation abundance and levels of methylation at the target sites of JmjC domain demethylases. Following succinate treatment, we observed a significant increase in di- and tri-methylation at the H3K9 residue, as well as a significant increase in H3K27me2 (**Fig. 4B**). Interestingly, these residues are among the targets of KDM4D and KDM6B, respectively, which we found to be the enzymes inhibited by the Ksu peptide in AlphaLISA assays (**Fig. 3C-D**). We also treated HepG2/C3A spheroids, which replicate more slowly than 2D cell culture. This is particularly relevant when assessing perturbations to silencing histone methylation marks that have turnover in the order of days, rather than hours like histone Kac^37,38^. In this culture system, 48 hours of succinate treatment increased levels of H3K9 di- and tri-methylation (**Fig. 4C**). After a week of succinate treatment, we also observed a significant increase in the levels of H3K27me3 and H3K36me3 marks (**Fig. 4C**).

**Figure 4.**
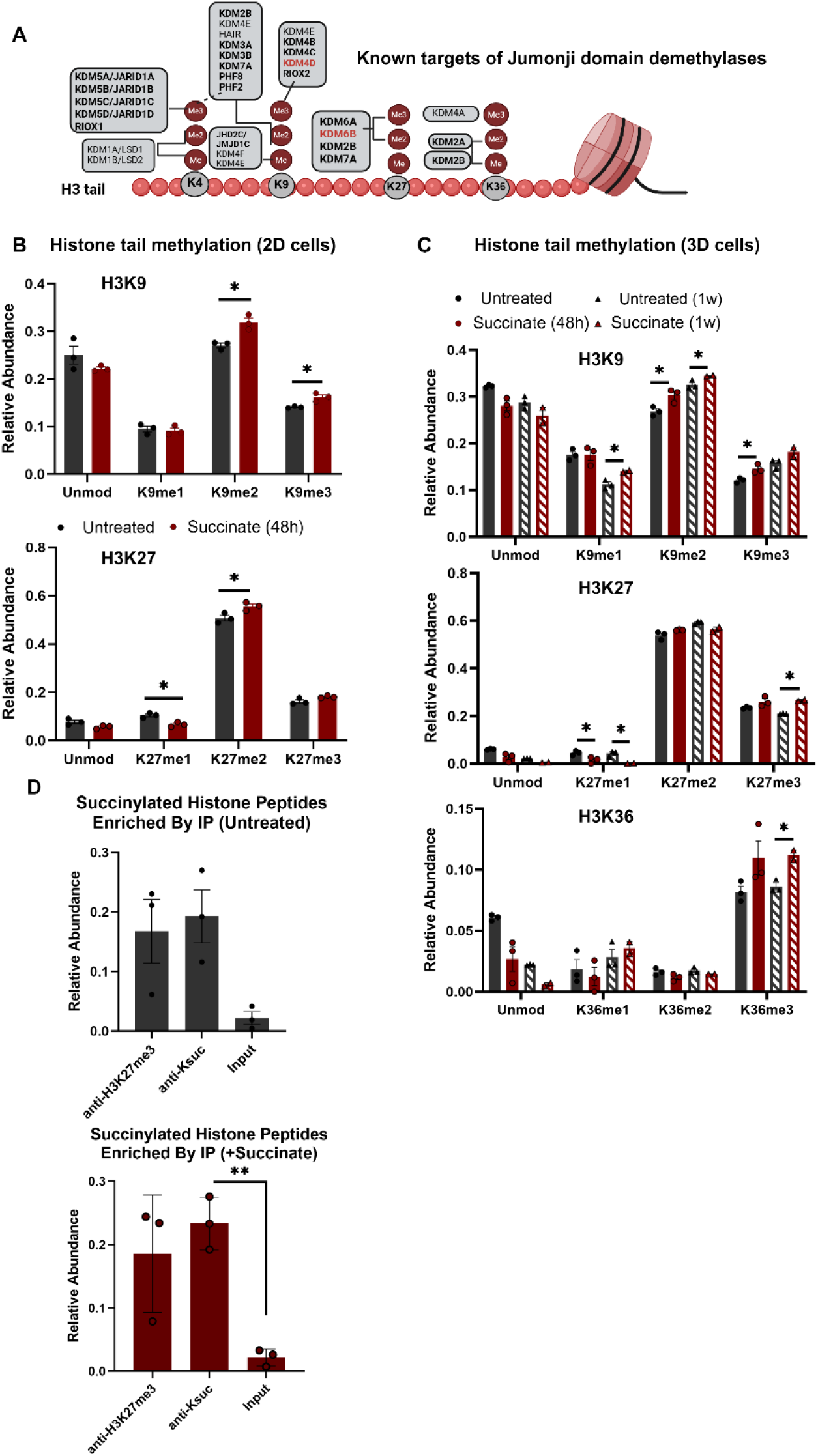
Increased global histone succinylation co-occurs with increased methylation at targets of Jumonji domain demethylases. (A) Known targets of JmjC domain demethylase family enzymes. (B) Relative abundance of methylation on targets of JmjC domain demethylases in HepG2/C3A cells treated with sodium succinate. (C) Relative abundance of methylation on targets of JmjC domain demethylases in HepG2/C3A 3D cells treated with sodium succinate. (D) ChIP-MS shows co-enrichment of succinylated histone peptides with H3K27me3 peptides. *Data are shown as mean ± SEM; *p-value <0*.*05, **p-value <0*.*01 in two-tailed Student’s t-test*.

To determine if histone succinylation coincides locally on chromatin with any changes in H3K9 or H3K27 methylation, we used chromatin immunoprecipitation followed by mass spectrometry (ChIP-MS) to assay whether nucleosomes modified with H3K27me3 also display enriched levels of histone succinylation (**Fig. 4D, top**). The converse approach proved to be impractical due to the weak binding of anti-succinyl antibodies (data not shown). We found that a subset of the nucleosomes immunoprecipitated with the anti-H3K27me3 antibody contained an enriched amount of Ksu marks. The experiment was repeated using HepG2/C3A cells treated with 10 mM sodium succinate and produced similar results (**Fig. 4D, bottom**). These results indicated that Ksu and H3K27me3 are enriched on the same nucleosomes (**Fig. S7E**).

### Transcriptomic analysis and chromatin profiling of succinate treated HepG2/C3A cells identifies candidate loci for co-occurring H3K27me3 and Ksu histone marks

To determine the transcriptional consequences of the observed increase in heterochromatic methylation on the histone H3 tail, we performed mRNA-seq on HepG2/C3A cells treated with 10 mM sodium succinate for 48 and 96 hours. Exogenous spike-in normalization was used to determine absolute mRNA fold changes^39^. We observed that transcriptional changes were minimal after 48 hours of treatment, but following 96 hours of treatment, approximately 20% of gene transcripts were significantly downregulated (**Fig. 5A-B**). As an additional control, we treated HepG2/C3A cells with 10mM sodium acetate to corroborate that the observed transcriptional effects were specific to succinate treatment. Acetate treatment initially increased transcriptional output (**Fig. S8C**), which is consistent with a global increase in histone acetylation. The transcriptional changes observed in the acetate-treated cells lacked enrichment in any ontology or pathway analysis and were distinct from the gene expression changes produced by succinate treatment (**Fig. S8E**). In contrast, pathway analysis of downregulated genes following succinate treatment identified downregulation of genes involved in DNA replication and translational machinery, including ribosome biogenesis, ribosomal subunits, and aminoacyl-tRNA biosynthesis (**Fig. S8F-G**). Consistent with these observations, JmjC demethylases have been implicated in the regulation of rDNA transcription, and their inhibition has been associated with reduced ribosome production^40–43^. However, these transcriptional changes cannot be directly attributed to histone succinylation alone without further context.

**Figure 5.**
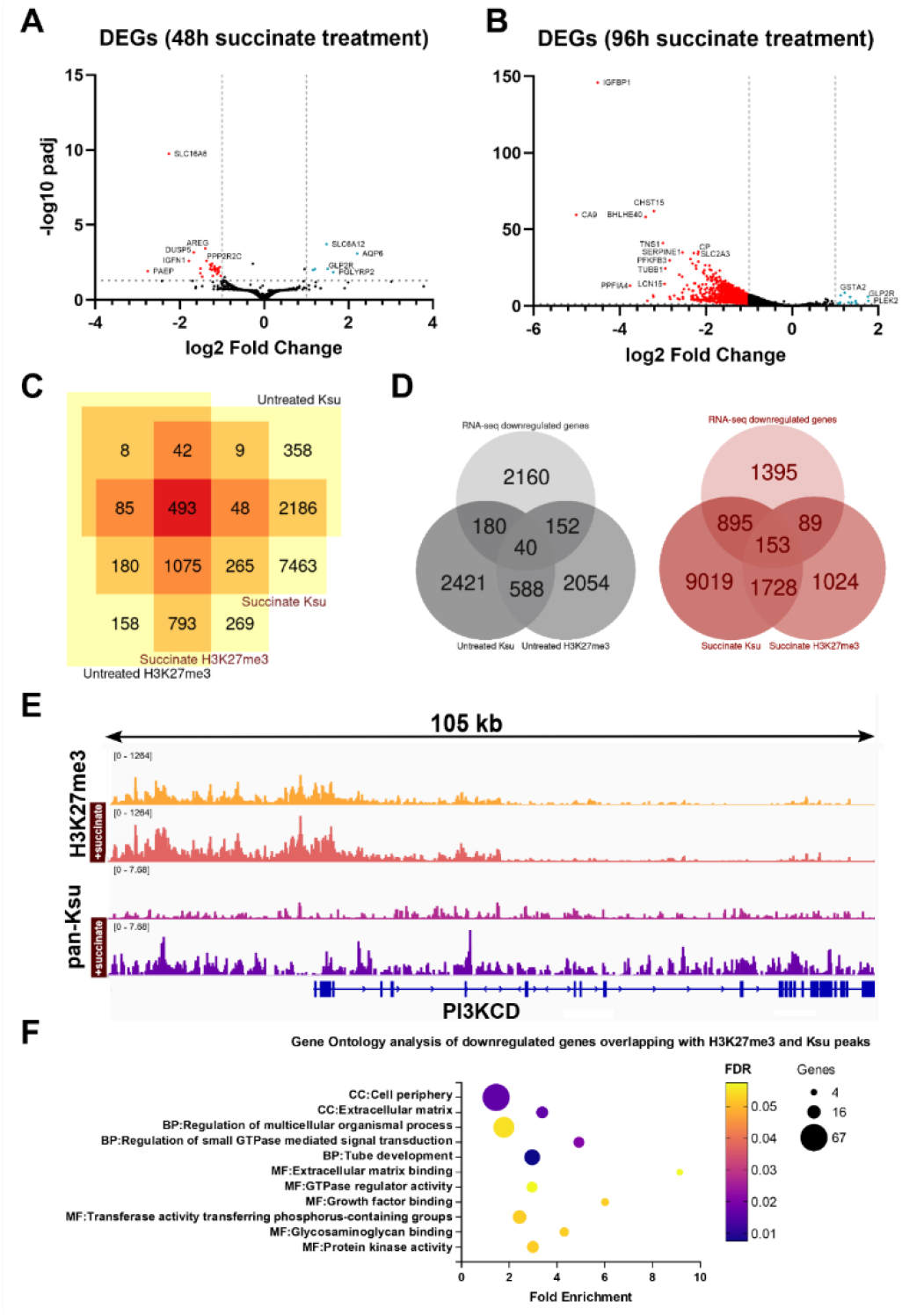
Transcriptomic analysis and chromatin profiling of succinate treated HepG2/C3A cells identifies candidate loci for co-occurring H3K27me3 and succinyl-lysine histone marks. Volcano plot of differentially expressed genes following (A) 48 hours or 96 hours of treatment with 10 mM sodium succinate using S2 spike-in normalization. (C) Venn diagram of overlap between genes with either H3K27me3 or Ksu peaks. (D) Venn diagram of genes with H3K27me3 or Ksu peaks (+succinate and untreated) and genes significantly downregulated in transcriptomic analysis following succinate treatment for 96 hours (FC threshold < -1, FDR < 0.05). (E) H3K27me3 and Ksu peaks at the PIK3CD gene locus. (F) Gene ontology analysis using iDEP^46^ of genes downregulated following 96 hours of succinate treatment, with co-occurring H3K27me3 and Ksu peaks at the gene locus (FC threshold < -1, FDR < 0.05).

Next, we performed a CUT&Tag protocol^44^ on HepG2/C3A cells treated with succinate, enriching for H3K27me3 and Ksu. The H3K27me3 and Ksu enriched regions showed partial overlap in selected gene loci, which we compared to the list of differently expressed genes from the RNA-seq experiment (**Fig. 5C-D**). We found that several genes which were significantly downregulated following succinate treatment (96 hours) had both H3K27me3 and Ksu peaks surrounding the transcription start site region of the gene. For example, the locus of *PIK3CD*, a gene encoding a subunit of a kinase involved in cell growth (P110δ), displayed ‘spreading’ of the characteristically broad H3K27me3 peak and a co-occurring Ksu peak (**Fig. 5E**). Another noteworthy downregulated gene displaying this pattern was hepatocyte nuclear factor-1 homeobox B (*HNF1B*) (**Fig. S8H**). *HNF1B* is a transcription factor involved in the development of several tissues and has recently been shown to have a role in mediating the proliferation and migration of hepatocellular carcinoma cells^45^.

Gene ontology analysis of the subset of genes that displayed both downregulated transcription and co-occurring H3K27me3 and Ksu peaks highlighted genes involved in kinase signaling and growth factor binding, reflecting reduced proliferation and metabolic modulation (**Fig. 5F**). These observations are consistent with the diminished growth phenotype of HepG2/C3A cells under long-term succinate treatment (**Fig. 1F**). Delayed transcriptional and phenotypic effects may be influenced by the slower kinetics of histone methylation^37^. Taken together, these results are consistent with a metabolite-transcription coupling model through which histone succinylation and H3K27me3 co-occur at a subset of gene loci and are a candidate crosstalk pattern for reduced transcriptional output.

## DISCUSSION

In this study, we demonstrate that despite the structural and chemical similarities to acetylation, histone succinylation recruits a unique set of chromatin modifying enzymes; the Jumonji domain demethylases.

We show that succinate treatment leads to elevated global histone succinylation (**Fig. 1A-B**), primarily accumulating on core histone lysine residues (**Fig. 1C-D**). Multiple evolutionarily unrelated JmjC domain demethylases can be pulled-down from the nuclear lysates of HepG2/C3A by a succinyl-lysine peptide pull-down (**Fig. 2E-F**).

Similarly, supraphysiological succinate treatment generated a chromatin fraction enriched with JmjC domain demethylases, compared to untreated cells (**Fig. 2I**). These observations led us to the finding that JmjC catalytic sites are structurally adept at binding Ksu peptides, which are predicted to occupy the αKG co-factor site by AlphaFold and Boltz2 models (**Fig. 2G-H**). The succinyl-lysine–JmjC interaction can be experimentally evidenced by succinyl-lysine peptide inhibition of demethylase assays, specifically KDM6B and KDM4D (**Fig. 3C-D**). In isolation from other binding determinants, the Ksu modification is a weak driver of affinity, but significantly stronger than Kac. We observed that succinate treatment corresponded with increased levels of methylation at the targets of JmjC domain demethylases (**Fig. 4B-C**), globally decreased transcriptional output (**Fig. 5B**), and downregulated pathways involved in cell proliferation (**Fig. 5F**). We used CUT&Tag chromatin profiling to identify several candidate gene loci with co-occurrence of lysine succinylation and H3K27me3 (**Fig. 5C**), along with decreased transcriptional output during succinate treatment (**Fig. 5D**).

A previous study on the putative functional roles of lysine succinylation found that elevated histone succinylation correlated with H3K4me3, and that promoter regions carrying both modifications had increased transcriptional output^9^. These observations are distinct but consistent with our own observations of the changes in methylation following succinate treatment (**Fig. 4**), suggesting a functional crosstalk between H3 tail lysine methylation and histone core succinylation. Histone PTM crosstalk is an epigenetic phenomenon where histone PTMs act in concert or in opposition to each other as part of their chromatin regulatory process^47–49^. Our proteomic assays have demonstrated that histone H3K64 is the most frequently succinylated residue for HepG2/C3A cells (**Fig. 1C**). H3K64 is outside the histone tail, where most histone PTMs having well characterized writer-reader-eraser pathways are located, including the target lysines of JmjC domain demethylases. However, both local and distal histone PTMs can display crosstalk (for example, H2BK123 ubiquitination maintaining H3K4 and K79 trimethylation)^50^. We propose a new function for histone succinylation, i.e. by inhibiting and sequestering JmjC domain demethylases. This enforces a dual-silencing mark with pre-existing H3 tail methylation. The previously observed relationship between H3K4me3 and Ksu suggests additional mechanisms of Ksu–JmjC–Kme3 crosstalk may be uncovered^9^.

Understanding the link between metabolism and chromatin regulation has become increasingly important in the study of aging and cancer. Recently, we reported preliminary evidence that histone succinylation may associate with longevity-related phenotypes, as individuals with projected longevity displayed elevated levels of this modification, and succinate supplementation improved strength and motor coordination in aged mice^51^. Given the well-established association between aging and heterochromatin loss^52^, our findings raise the possibility that histone succinylation may contribute to the maintenance of repressive chromatin states through inhibition of JmjC domain demethylases. Several cancers are driven by mutations in succinate dehydrogenase (SDH), causing an increase in succinate that leads to product inhibition of αKG-dependent dioxygenases, including JmjC demethylases^24^. Histone lysine succinylation has also been shown to have a role in several cancers; for example, succinylation of H3K79 activates the transcription of oncogenic genes and promotes cell proliferation in glioblastoma and pancreatic carcinoma^12,53^. It is therefore possible that our supraphysiological succinate treatment may be directly responsible for a portion of the observed increase in methylation in our data. Conversely, this also suggests that the observed JmjC demethylase inhibition and elevated methylation in an SDH-deficient model is multifactorial, driven by both elevated succinate and histone succinylation.

The model we have presented here describes a distinct relationship between histone succinylation and an abundant class of αKG and iron-dependent catalytic sites, fitting in an additional piece of the puzzle of uncharacterized rare histone PTMs. This study lays the groundwork for future research into the functional consequences of the inhibition of Jumonji domain demethylases by histone succinylation and its implications for the modulation of metabolites for therapeutic purposes.

## RESOURCE AVAILABILITY

### Materials availability

This study did not generate new unique reagents.

### Data and code availability

RNA-seq and CUT&Tag data have been deposited at GEO GSE331257 and GSE331259 and are publicly available.

The proteomics data are available in ProteomeXchange (PRIDE) under the project number PXD078425. Reviewers can access the data using the token “WCjorsrbofFh”. Alternatively, reviewers can access the dataset by logging in to the PRIDE website using the following account details: Username; Password: HVVpZZdr5wly.

## ACKNOWLEDGMENTS

The Sidoli lab gratefully acknowledges funding from the Aging Biology Foundation, the Hevolution Foundation (AFAR), the ERCM-CFAR Center for AIDS Research, the NIAID for the PROVIDENT grant (1U19AI181977), the Einstein-Mount Sinai Diabetes center, and the NIH Office of the Director (S10OD030286). NWS was supported by R35GM156596 and AC was supported by T32HL091804. DS was supported by R01GM108646 and Hevolution HF-GRO-23-1199128-32 and IK was supported by Hevolution HF-GRO-23-1199128-32 and T32GM149364. We thank Dr. Radha Charan Dash for additional JmjC models, and colleagues from Reaction Biology Corporation for insightful assay controls.

## AUTHOR CONTRIBUTIONS

Conceptualization, S.G., J.P.M., and Si.S.; methodology, S.G., J.P.M., S.S., A.P., K.D., A.C., N.W.S., and Si.S.; investigation, S.G., S.S., I.K., K.D., A.P., A.C., and N.W.S.; formal analysis, S.G., A.P., A.C., N.W.S., J.P.M., and Si.S.; resources, D.S., J.P.M., N.W.S., and Si.S.; writing – original draft, S.G.; writing – review & editing, S.G., J.P.M., D.S., and Si.S.; supervision, D.S., J.P.M., and Si.S.; funding acquisition, J.P.M. and Si.S.

## SUPPLEMENTAL INFORMATION

**Document S1. Figures S1–S6, Tables S1-4**

**Document S2: Key for experimental raw files**

**Document S3: DIA-NN search results peptide pull-down**

**Document S4: Proteome Discoverer search results succinate treated nuclear proteomics**

## METHOD DETAILS

### Flat cell culture

The human hepatocellular carcinoma HepG2/C3A cell line was obtained from the American Type Culture Collection (ATCC, CRL-10741). The cells were cultured in Dulbecco’s Modified Eagle’s Medium (DMEM, containing 4.5 g/L glucose) supplemented with 10% Fetal Bovine Serum (FBS), 1% GlutaMAX, 1% NEAA, and 0.5% Penicillin/Streptomycin (all Corning). Cells were kept in a controlled atmosphere (5% CO2 incubator at 37 °C). Cells were subcultured by trypsinization with 0.05% Trypsin-EDTA diluted 1:1 in Hanks balanced salt solution (Corning) for 5 minutes, and centrifuged for 5 minutes at 140 g. The number of cells was estimated using the Corning Cell Counter and CytoSMART Cloud App (Corning).

### Spheroid culture

HepG2/C3A spheroids were cultured according to the protocol in^54^. In brief, HepG2/C3A spheroids were formed using a cell suspension (1.2 × 10^6^ cells diluted in growth media) added to an ultra-low attachment 24-round bottom plate containing microwells (Elplasia®, Corning). The cells were incubated overnight to aggregate into spheroids and then transferred into a prepared ClinoReactor with fresh growth media. Growth media was changed three times a week, and rotational speed adjusted with spheroid growth. Spheroids were maintained in fully supplemented growth media for at least 35 days prior to treatment.

### Histone extraction from sodium succinate treated HepG2/C3A cells

In order to elevate the levels of histone succinylation, cells were treated with sodium succinate. Sodium succinate (S9637) was dissolved in water and sterile filtered, then added to complete growth media minus non-essential amino acids to a final concentration of 10 mM. HepG2/C3A flat cells were treated with 10 mM sodium succinate for 48 hours, and then trypsinized using 0.5% Trypsin/EDTA diluted 1:1 in Hanks solution (Corning) for 5 min, followed by centrifugation at 140 g for 5 min. The cell pellet was then resuspended in Hanks solution, followed by centrifugation at 140 g for 5 min. The supernatant was removed and the cell pellets were stored in -80C prior to histone extraction. The media for HepG2/C3A spheroids (>35 days) was exchanged to one containing 10 mM sodium succinate, and fresh media containing 10 mM sodium succinate was supplied every 48 hours. Spheroids were collected at 48 hour and 1 week time points, washed with Hanks solution, and centrifuged at 1,000 rpm for 5 min. The supernatant was discarded and dry pellets were stored at -80 C.

Histone proteins were extracted from cell pellets, as described by^54^. Briefly, histones were acid extracted with cold 0.2 M sulfuric acid (5:1, sulfuric acid:pellet) and incubated for 2 h at 4ºC with constant rotation. Histones were then precipitated with 33% trichloroacetic acid (TCA) overnight at 4ºC. The supernatant was removed, tubes were washed with ice-cold acetone containing 0.1% HCl, centrifuged and washed again with 100% ice-cold acetone. The supernatant was discarded, and the pellet was dried using the SpeedVac. Before derivatization, histones were resuspended in 50 mM ammonium bicarbonate (pH 8) containing 20% acetonitrile. In the fume hood, 2 µl of propionic anhydride and 10 µl of ammonium hydroxide (all Sigma Aldrich) were added to the samples. The mixture was incubated for 5 min and the procedure was repeated. Histones were then digested with 1 µg of sequencing grade trypsin (Promega) (1:20, enzyme:sample) diluted in 50 mM ammonium bicarbonate, pH 8, and incubated overnight at 37ºC. The derivatization reaction was repeated to derivatize peptide N-termini. The peptides were dried in the Speedvac and stored in -80ºC.

### Peptide pull down assay

50 ul of streptavidin beads (item number) per sample were washed three times with 1 mL peptide binding buffer (150 mm NaCl, 50 mM Tris, 0.1% NP-40, pH 8). 25ul of each synthetic peptide (1 mg/mL) diluted in 475 ul peptide binding buffer were incubated with streptavidin beads rotating at 4C overnight. Prior to protein lysate loading, beads conjugated to peptides were washed three times with 1 mL peptide binding buffer. The nuclear proteome was isolated from HepG2/C3A cell pellets (2×10^6) grown in flat culture. Briefly, the cell pellet was resuspended in five volumes of cold buffer A (10 mM ammonium bicarbonate pH 8, 1.5 mM MgCl2, and 10 mM KCl), incubated on ice for 10 min and centrifuged for 5 min at 400 g. The supernatant was removed and the pellet was resuspended in two volumes of buffer A containing 0.15% NP-40 and protease inhibitors and then centrifuged for 15 min at 3,200 g. The pellet was washed with ten volumes of PBS, centrifuged for 5 min at 3,200 g, and the supernatant was discarded. The pellet was resuspended with two volumes of buffer C (420 mM NaCl, 20 mM ammonium bicarbonate pH 8.0, 20% glycerol, 2 mM MgCl2, 0.2 mM EDTA, 0.1% NP-40, 10 mM sodium butyrate, 0.5 mM DTT, and complete protease inhibitors) and incubated for 1 h at 4ºC, on a rotating wheel. The suspension was centrifuged for 45 min at 20,800 g at 4ºC. The supernatant contained the soluble nuclear fraction, which was diluted to 500 uL in protein binding buffer ((150 mm NaCl, 50 mM Tris, 10 uM ZnCl2, 0.5 mM DTT, 1:100 protease inhibitors (PI78429), 10 mM sodium butyrate, pH 8), then incubated with the biotinylated histone peptides conjugated to the streptavidin beads at 4C, rotating, overnight. Streptavidin beads alone were used as a control to remove non-specific interaction between proteins in the lysate and the beads. Following the incubation, the beads were transferred to an Orochem OF1100 96-well plate and washed 3 times with protein binding wash buffer (350 mM NaCl, 50 mM Tris, 10 uM ZnCl2, pH 8) to remove non-specific binding. Proteins were reduced with 100 ul of 5 mM DTT in 50 mM ammonium bicarbonate for 1 hour at room temperature, then alkylated with 100 uL of 20 mM iodoacetamide in 50 mM ammonium bicarbonate for 30 minutes in the dark. The solution was removed via vacuum prior to trypsin digestion. The proteins were then digested off the beads using 500 ng trypsin in 25 uL ammonium bicarbonate, eluted with 25 uL of 60% acetonitrile in 0.1% TFA, and dried prior to desalting.

### Desalting

Samples were desalted prior to mass spectrometry (MS) analysis. Briefly, samples were resuspended in 100 µl of 0.1% TFA and loaded into a 96-well filter plate (Orochem) packed with Oasis HLB C-18 resin (Waters), which was equilibrated using 100 µl of the same buffer. Samples were washed with 100 µl of 0.1% TFA, eluted with 70 µl of a buffer containing 60% acetonitrile and 0.1% TFA, and dried using SpeedVac.

### Histone post-translational modification (PTM) analysis

After desalting, samples were resuspended in 10 µl of 0.1% TFA and loaded onto a Dionex RSLC Ultimate 300, coupled online with an Orbitrap Fusion Lumos (all Thermo Scientific). Chromatographic separation was performed using a two-column system, consisting of a C-18 trap cartridge (300 µm ID, 5 mm length) and an analytical column (75 µm ID, 25 cm length) packed in-house with reversed-phase Repro-Sil Pur C18-AQ 3 µm resin. Histone peptides were separated using a 45 min gradient from 4–30% buffer B (buffer A: 0.1% formic acid, buffer B: 80% acetonitrile + 0.1% formic acid) at a flow rate of 300 nl/min. The mass spectrometer was set to acquire spectra in a data-independent acquisition (DIA) mode. The full MS scan was set to 300–1100 m/z in the orbitrap with a resolution of 120,000 (at 200 m/z) and an AGC target of 5 × 105. MS/MS was performed in the orbitrap with sequential isolation windows of 50 m/z with an AGC target of 2 × 105 and an HCD collision energy of 30.

Raw files were imported into EpiProfile 2.0 software^26^. From the extracted ion chromatogram, the area under the curve was obtained and used to estimate the abundance of each peptide. To achieve the relative abundance of PTMs, the sum of all different modified forms of a histone peptide was considered 100% and the area of the particular peptide was divided by the total area for that histone peptide in all of its modified forms. The relative ratio of two isobaric forms was estimated by averaging the ratio for each fragment ion with different mass between the two species. The resulting peptide lists generated by EpiProfile were exported to Microsoft Excel and further processed for a detailed statistical analysis.

### Proteomic analysis

After desalting, samples were resuspended in 10 µl of 0.1% TFA and loaded onto a Dionex RSLC Ultimate 300, coupled online with an Orbitrap Fusion Lumos (all Thermo Scientific). Chromatographic separation was performed using a two-column system, consisting of a C-18 trap cartridge (300 µm ID, 5 mm length) and an analytical column (75 µm ID, 25 cm length) packed in-house with reversed-phase Repro-Sil Pur C18-AQ 3 µm resin. Samples were separated using a 60 or 90 min gradient from 4 to 30% buffer B (buffer A: 0.1% formic acid, buffer B: 80% acetonitrile + 0.1% formic acid) at a flow rate of 300 nl/min.

Peptide pull-down: The mass spectrometer was set to acquire spectra in the data-independent acquisition (DIA mode).

Nuclear proteomics: The mass spectrometer was set to acquire spectra in a data-dependent acquisition (DDA) mode. The full MS scan was set to 300–1200 m/z in the orbitrap with a resolution of 120,000 (at 200 m/z) and an AGC target of 5 × 105. MS/MS was performed in the ion trap using the top speed mode (2 s), and AGC target of 1 × 104 and an HCD collision energy of 35.

The DDA MS raw data were processed using Proteome Discoverer software (v2.5, Thermo Scientific), SEQUEST search engine, and the SwissProt human database (updated June 2021). The DIA MS raw data was processed using DIA-NN (v1.8, https://github.com/vdemichev/diann) using the SwissProt human database (updated June 2021). Both searches included variable modification of N-terminal acetylation and fixed modification of carbamidomethyl cysteine. Trypsin was specified as the digestive enzyme with two missed cleavages allowed. Mass tolerance was set to 10 ppm for precursor ions and 0.2 Da for the product ions. Peptide and protein false discovery rate was set to 1%. Prior to statistical analysis, proteins were log2 transformed, normalized by the average value of each sample and missing values were imputed using a normal distribution 2 standard deviations lower than the mean as described^55^. Statistical regulation was assessed using heteroscedastic T-test (if p-value < 0.05). Data distribution was assumed to be normal, but this was not formally tested.

### SILEC-SF labeling and metabolomics

HepG2/C3A spheroids were cultured and treated with 10 mM sodium succinate as previously described. In parallel, according to the protocol described in^30^, HepG2 cells were cultured for >9 generations in DMEM containing ^13^C_3_^15^N_1_-pantothenate and minimal unlabeled pantothenate to generate cells with labeled acyl-CoAs highly enriched for ^13^C_3_^15^N_1_-pantothenate (VB5* cells). Before fractionation, These VB5* HepG2 cells were added 1:1 to the treated HepG2 spheroids and lysed (Buffer: 250 mM sucrose, 15 mM Tris–HCl (pH 7.5), 60 mM KCl, 15 mM NaCl, 5 mM MgCl2, 1 mM CaCl2, 0.1%NP-40 adjusted to pH 7.4). Nuclei were pelleted by centrifugation at 600 g for 5 min at 4C. The supernatant (non-nuclear fraction) was quenched in 10% (w/v) trichloroacetic acid. The nuclear pellet was washed with lysis buffer without NP-40 and then quenched in 10% (w/v) trichloroacetic acid. Samples were stored at -80C before processing. Acyl-CoAs were measured by liquid chromatography-high resolution mass spectrometry as previously described in detail^56^. Briefly, samples were quenched in 1 mL 10% (w/v) trichloroacetic acid in water for extraction. Samples were homogenized by using a probe tip sonicator in 0.5 second pulses 30 times then centrifuged at 17,000 x g for 10 min at 4°C. Supernatant was purified by solid phase extraction (SPE) cartridges (Oasis HLB 10 mg, Waters) that were conditioned with 1 mL of methanol and 1 mL of water. Acid-extracted supernatants were loaded onto the cartridges and washed with 1 mL of water. Acyl-CoAs were eluted with 1 mL of 25 mM ammonium acetate in methanol and evaporated to dryness under nitrogen. Samples were resuspended in 50 µL of 5% (w/v) 5-Sulfosalicylic acid and 20 µL injections were analyzed on an Ultimate 3000 UHPLC using a Waters HSS T3 2.1×100mm 3.5 µm column coupled to a Q Exactive Plus with a HESI II probe operating in positive ion mode. The analysts were blinded to sample identity during processing and quantification. Instruments were controlled via XCalibur 4.1 and data was analyzed on Tracefinder 5.1 using a 5ppm window from the predominant ion positive. Area under the curves for each analyte was normalized to the matched internal standard or the nearest surrogate internal standard.

### Boltz2 Modelling

The JmjC domain FASTA sequences were pulled from the Uniprot database^57^. Boltz2 modelling of different KDM enzymes was conducted using the following inputs in the Boltz2 structure prediction algorithm in the Tamarind.bio server^58^. All inputs remained the same for modelling all JmjC enzymes in the modelling analysis, swapping the FASTA sequence of the JmjC domain (sequence number used for modelling of each enzymes can be found in Table S2). Ten output models were selected for each job. The models were downloaded as PDB files and analyzed in PyMol^59^. Additional information about the inputs can be found in Supplementary Document S1.

### Expression and purification of KDM6B

The JMJD3 plasmid was a gift from Nicola Burgess-Brown (Addgene plasmid # 39069 ; http://n2t.net/addgene:39069 ; RRID:Addgene_39069). Expression vectors were transformed into Rosetta 2(DE3) competent cells, which were expanded into a starter culture overnight at 37C. Sterilized Luria (Miller) broth supplemented with 50 μ/mL kanamycin and 25 μ/mL chloramphenicol was inoculated with starter culture and incubated with rotation at 37C for several hours. Once an optical density (O.D.) of 0.6 was reached, cultures were cooled at 18C for 1 hour and induced with 0.25 mM IPTG, then incubated with rotation overnight (12-16 hours) at 18C. The bacteria were pelleted by centrifugation, resuspended in ice cold lysis buffer (100mM Tris-HCl pH 8.0, 300 mM NaCl, 10% glycerol), and lysed by Emulsiflex (15,000-18,000 psi). KDM6B was initially purified with Ni-NTA resin (Thermo). Briefly, pre-equilibrated Ni-NTA resin was incubated with lysate at 4C with gentle rotation for 1 hour. Beads were washed with 1.5 column volumes (CV) of lysis buffer supplemented with 30 mM imidazole. Protein was eluted three times with 1.5 CV of lysis buffer supplemented with 250 mM imidazole. The first and second elutions were combined and volume was reduced with an Amicon spin filter. KDM6B was purified further with size exclusion chromatography with a Superdex 200 10/300 column (20 mM Tris pH 8, 150 mm NaCl, 10% glycerol, 1mM DTT, 0.5 mM EDTA). The protein was then buffer exchanged into a low salt buffer (20 mM Tris pH 8, 50 mm NaCl, 10% glycerol, 1mM DTT, 0.5 mM EDTA) and purified further with MonoQ ion exchange chromatography. Purified KDM6B was stored in 50% ion exchange buffer/50% glycerol).

### Surface Plasmon Resonance (SPR)

The interaction between the KDM6B protein and the two peptides (Biotinylated acetyl-lysine and succinyl-lysine histone H3K64 peptides (Table S1) ) was analyzed by SPR on a Biacore 1S+ (Cytiva) system using Series S Biotin CAPture kit (Cytiva) at 25 °C, with 10 mM HEPES pH 7.4, 150 mM NaCl, 2% DMSO, 1 mM a-KG, 100 µM NiCl2, 100 µM ascorbic acid as the running buffer. Biotin CAPture reagent (diluted 1:4 in running buffer) was first immobilized on the chip surface following manufacturer’s protocol. The H3K64 peptides were captured as ligands (∼1000 response units) on parallel channels by running 40 µg/mL peptide solution in running buffer. Data was acquired by injecting 1 and 5µM KDM6B in running buffer as analytes in separate cycles at 30 µL/min for 30 s to observe association, followed by dissociation in running buffer for 30 s. Double referencing (by subtracting responses from a blank cycle and a channel immobilized with CAPture reagent alone) was performed to exclude response from bulk effects and non-specific interaction of the analyte to chip surface, respectively. The chip was regenerated as per the manufacturer’s protocol between replicate concentration series. All data analyses were performed using the GraphPad Prism software. The dissociation constants (Kd) of the enzyme: peptide interaction could not be calculated.

### Western blot validation of antibodies

Histones were isolated from HepG2/C3A cells as described previously, and HepG2 cell lysates were produced using the nuclear fractionation protocol described previously. Samples were boiled in the presence of BME and 40 ug of cell lysate and ∼5 ug of histones were loaded on a 4-20% SDS-PAGE gel. Proteins were transferred to PVDF membrane overnight at 24v and blocked for 1 hour at room temperature with blocking buffer: 5% milk in TBS-T 0.1% for anti-H3K27me3, anti-H3, and anti-Ksu (cell signaling), and 3% milk in TBS-T 0.1% for Invitrogen Ksu antibody (PA5-120740). Membranes were incubated with primary antibodies in blocking buffer overnight at 4C (1:500 for anti-Ksu, 1:1000 for anti-H3 and H3K27me3), followed by HRP-conjugated secondary antibodies (1:10,000). Signals were detected using ECL substrate on a BioRad imaging system.

### ChIP-MS

The chromatin immunoprecipitation protocol was adapted from^60^. Briefly, 20 ul of Protein A/G Dynabeads™ beads per sample (Thermo) were washed with 0.5% BSA in PBS three times, then incubated with primary antibodies overnight at 4C (5 ug anti-Ksu, 1 ug anti-H3K27me3). HepG2 cells were treated with sodium succinate as previously described. 4×10^6 cells per replicate were lysed (50 mM HEPES, 85 mM KCl, 0.5% NP-40, protease inhibitors), incubated on a nutator at 4C, then centrifuged for 10 min at 400 x g, 4C. Supernatant was discarded and nuclei were resuspended in a second lysis buffer (10 mM Tris pH 8, 150 mM NaCl, 1 mM EDTA, 1% NP-40, 0.1% SDS, 0.1% Na-deox) and sonicated in a Diagenode Pico sonicator for 16 cycles of 30 s on, 30 s off. Samples were then cleared by centrifugation at 16,000 x g, 4° C, 10 min. 50 ul of supernatant was kept as input and the remainder was incubated with the antibody-coupled beads overnight at 4C. Beads were then washed 3 times each with three wash buffers: 1; 20 mM Tris pH 8, 150 mM NaCl, 0.1% NP-40, 2; 20 mM Tris pH 8, 500 mM NaCl, 3; 10 mM Tris pH 8, 250 mM LiCl, and a final wash with 20 mM Tris. Derivatization and digestion of histone proteins bound to antibody-coupled beads was performed as previously described, and histone proteins were digested off the beads, which were removed during the desalting step.

### RNA-seq

HepG2 cells were treated with sodium succinate or sodium acetate for 48 and 96 hours as previously described. *Drosophila* S2 cells were cultured with S2 media, supplemented with 10% FBS and 1% PennStrep. 2.5×10^5 S2 cells were added for each 1×10^6 HepG2 cells, and were frozen in DNA/RNA Shield prior to RNA extraction. Total RNA was extracted with the Direct-zol RNA Miniprep Kit (Zymo R2050). Library preparation was done by Novogene Genetics US and sequencing was done on an Illumina platform using 150 nt paired-end libraries with 30 million reads (9Gb data per sample), with 4 independent biological replicates per condition. RNA-seq reads were trimmed and aligned to a composite human (hg38)-*Drosophila melanogaster* (dm6) genome using the Nextflow nf-core RNAseq pipeline (v3.21.0), with processes containerized using Singularity^61–63^. Differential abundance analysis was performed using DESeq2^64^ with a custom script (https://github.com/Shechterlab/DESeq2_with_SpikeInNormalization). Gene ontology and pathway analysis of differentially expressed genes was done with the iDEP (v2.01) web application^46^. Venn diagrams were made using the Intervene Shiny app and edited further with BioRender^65^.

### CUT&Tag

HepG2/C3A spheroids were treated with sodium succinate for 48 hours as previously described. The equivalent of 300,000 cells was used for each reaction. The iDeal CUT&Tag kit for Histones (Diagenode #C01070020) was used for the reaction, with 1 ug and 0.5 ug of Ksu antibody and H3K27me3 antibodies used respectively. Pooled and barcoded libraries were sequenced at Novogene Genetics US on an Illumina platform using 150 nt paired-end sequencing on a 10B flow cell (375Gb output per lane). Reads were trimmed and aligned and peak calling was done using the nf-core cutandrun pipeline (v3.2.2), using the SEACR peak caller, with processes containerized using Singularity^61–63,66^. Annotation of called peaks was performed using the ChIPseeker package^67,68^.

