## Supplementary material for "Histone succinylation directly inhibits Jumonji domain demethylases and stabilizes repressive chromatin states": Document S1

<u>**Supplementary Information**</u>

**This PDF includes:**

Supplementary Figure 1 .......... 2
Supplementary Table 1 .......... 3
Supplementary Table 2 .......... 3
Supplementary Table 3 .......... 4
Supplementary Figure 2 .......... 4
Supplementary Figure 3 .......... 5
Supplementary Table 4 .......... 5
Supplementary Figure 4 .......... 6
Supplementary Figure 5 .......... 7
Supplementary Figure 6 .......... 8
Supplementary Figure 7 .......... 9
Supplementary Figure 8 .......... 10
Supplementary Information for Boltz2 .......... 11

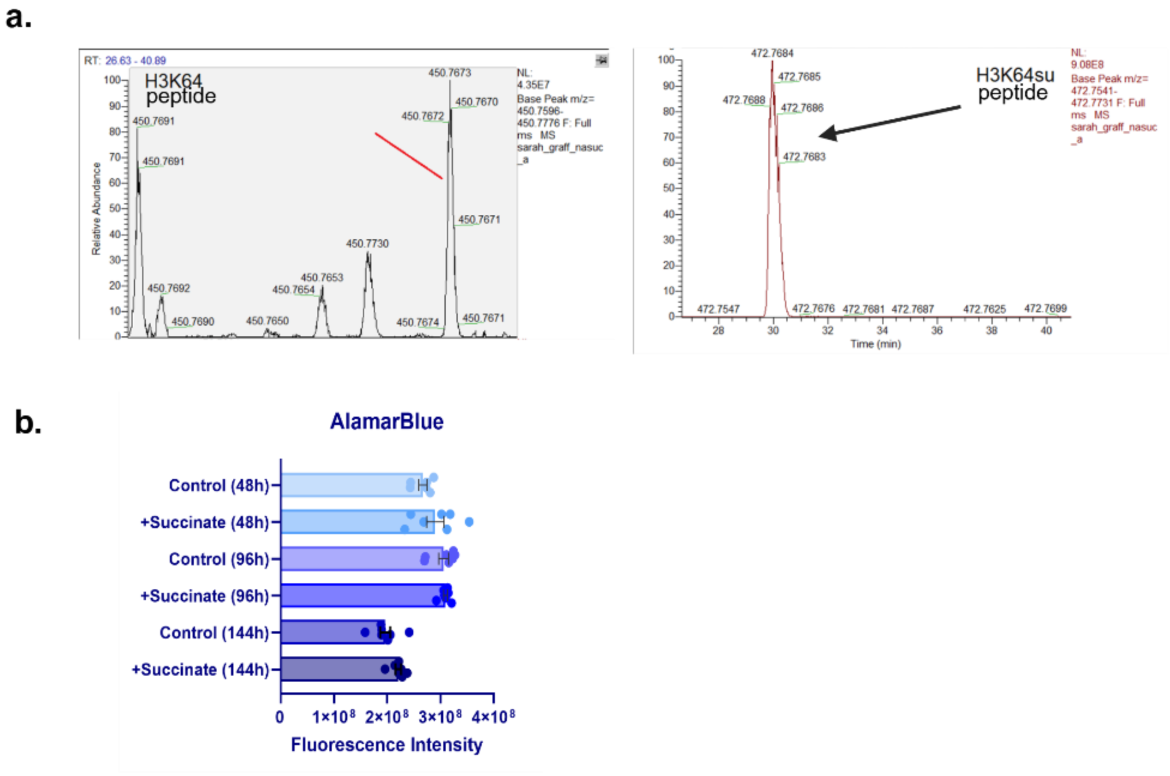


**Supplementary Fig. S1**. (a) Chromatogram of unmodified and succinylated histone H3K64-containing synthetic peptide. (b) AlamarBlue assay of HepG2/C3A cells treated with 10 mM sodium succinate for 48, 96, and 144h. No significant decrease in viability was observed.

**Table S1**: Synthetic peptides for proteomic and biochemistry assays

| Peptide sequence | Assay |
|---|---|
| AIRRYQKSTELLIRKIRRYQKSTEGK(Biotin) | Peptide pull-down |
| AIRRYQKSTELLIRK(Succinyl)IRRYQKSTEGK(Biotin) | Peptide pull-down/SPR |
| AIRRYQKSTELLIRK(Acetyl)IRRYQKSTEGK(Biotin) | Peptide pull-down/SPR |
| ATKAAR-Kme3-SAPSTGGVKKPHRYRPGGGK(Biotin)-NH2 | Succinate-Glo |
| AARK(Succinyl)SA-NH2 | AlphaLISA |
| AARK(Acetyl)SA-NH2 | AlphaLISA |
| AARK(Maleyl)SA-NH2 | AlphaLISA |
| AARK(Malonyl)SA-NH2 | AlphaLISA |
| AARK(Fumaryl)SA-NH2 | AlphaLISA |
| AARK(Glutaryl)SA-NH2 | AlphaLISA |

**Table S2**. Interactions observed in the KDM–KSu peptide models.

| Protein | Boltz2 Score (ipTM) | Average Cα RMSD (Å) *vs.* ref. | RMSD Referenc e PDB id | Residues modelled | Register of KSu in active site | Register reference PDB id |
|---|---|---|---|---|---|---|
| **KDM6B** | 0.966 | 0.43 | 6F6D | 1141-1643 | plus 1 | 4EZH |
| **KDM4A** | 0.943 | 0.18 | 2P5B | 2-350 | plus 1 | 2OQ6 |
| **KDM4C** | 0.985 | 0.32 | 4XDO | 10-347 | (plus 1)* | 4HON |
| **KDM4D** | 0.957 | 0.25 | 4HON | 12-341 | plus 1 | 4HON |
| **KDM5A** | 0.917 | 0.25 | 5IVB | 1-588 | (plus 1)* | 4HON |
| **KDM5B** | 0.946 | 0.27 | 5A1F | 26-770 | (plus 1)* | 4HON |
| **KDM5C** | 0.957 | 0.63 | 5FWJ | 468-634 | (plus 1)* | 4HON |
| **KDM3A** | 0.924 | 0.60 | (4C8D)* | 1050-1290 | (plus 1)* | 4HON |
| **KDM3B** | 0.929 | 0.38 | 5RAW | 1380-1728 | (plus 1)* | 4HON |
| **KDM2A** | 0.933 | 0.24 | 2YU1 | 1-517 | inverted | 4QX7 |
| **KDM2B** | 0.957 | 0.40 | (4QX7)* | 160-360 | (inverted)* | 4QX7 |
| **PHF8** | 0.93 | 0.40 | 3KV4 | 37-483 | plus 1 | 3KV4 |

(*) The crystal structure of a evolutionarily related KDM was used for the RMSD calculation and/or the register reference. ipTM output in Boltz2 stands for Interface Predicted Template Modeling score and is a key confidence metric used to rank the reliability of docked target-ligand structures. Values above 0.8 generally indicate a highly confident interface prediction. RMSD Reference PDB was the PDB structure that was used to compare the modelling outputs against a PDB structure of the protein. The register reference PDB ID was the PDB used to compare the positioning of the arginine and lysine residues of the modelled KSuc peptide to the known binding position of lysine containing peptides binding to the KDM proteins.

**Table S3**. Interactions predicted in the KDM–KSu peptide models.

| Protein | Pose A count (Fe❧O=C interaction)[a] | Pose B count (Fe–$O_2C$ interaction)[a] | Protein Lys in pose A salt bridge[b] | Distance Fe–Lys (Ne) in active site[c] | Count of poses with Arg bound in site | Inhibition in assays ($EC_{50}$ < 200 μM) |
|---|---|---|---|---|---|---|
| **KDM6B** | 10 | – | K1381 | 10.0 Å | 3 | Yes |
| **KDM4A** | 3 | 7 | K206 | 10.0 Å | 10 | No |
| **KDM4C** | 5 | 5 | K208 | 9.6 Å | 10 | No |
| **KDM4D** | 10 | – | K210 | 9.9 Å | 10 | Yes |
| **KDM5A** | 10 | – | K501 | 10.3 Å | – | No |
| **KDM5B** | 7 | – | K517 | 10.3 Å | – | No |
| **KDM5C** | 9 | – | K532 | 10.4 Å | – | No |
| **KDM3A** | 10 | – | K212 | 10.2 Å | 10 | Yes |
| **KDM3B** | 10 | – | K336 | 10.0 Å | – | Yes |
| **KDM2A** | 10 | – | K229 | 10.1 Å | – | No |
| **KDM2B** | 10 | – | K101 | 10.4 Å | 10 | No |
| **PHF8** | 9 | – | K264 | 9.6 Å | 1 | n/a |

[a] Pose A and pose B are shown in Fig. S2. [b]The JmjC catalytic site has a lysine within the αKG pocket, which forms a salt bridge in Pose A. [c]The distance spanned by the Ksu group in the αKG pocket can be expressed by the distance from the pocket lysine to the iron co-factor. Demethylase assays inhibited >50% by the peptide AARK(su)SA at a concentration of 250 μM or lower $IC_{50}$ (see Fig. 2).

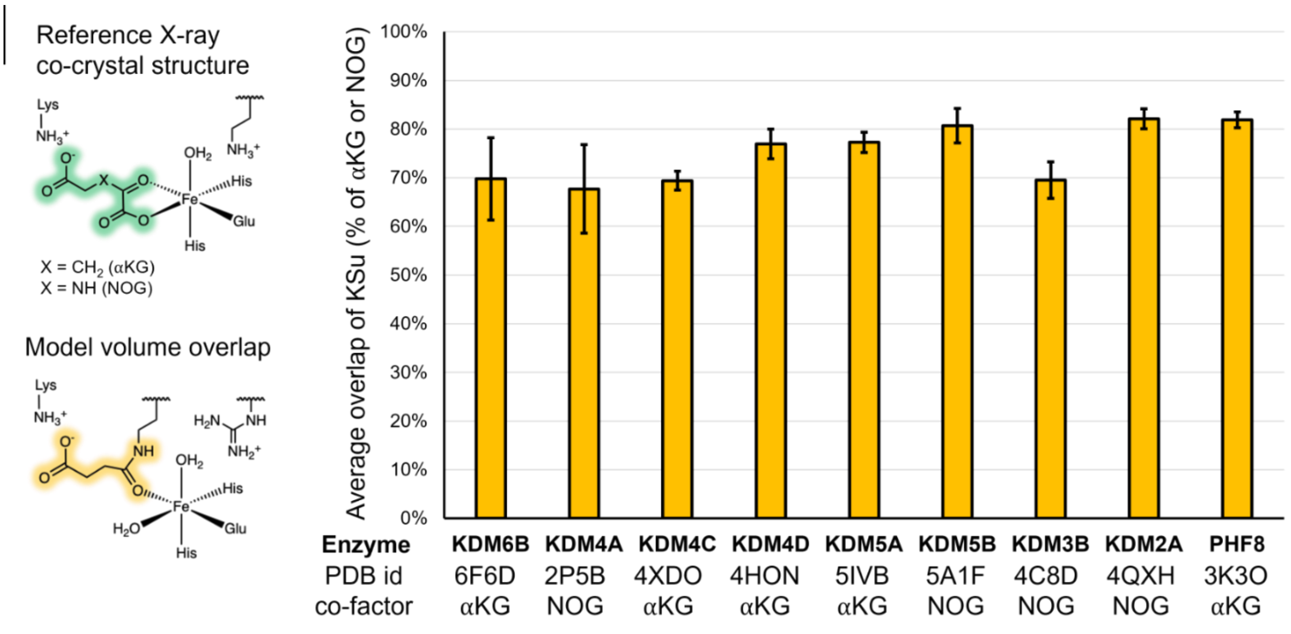


**Supplementary Figure S2**. Occupancy of the αKG pocket as calculated by the superimposed volume of JmjC demethylases co-crystallized with αKG or NOG and the KDM–KSu peptide models in Supplementary Table S3.

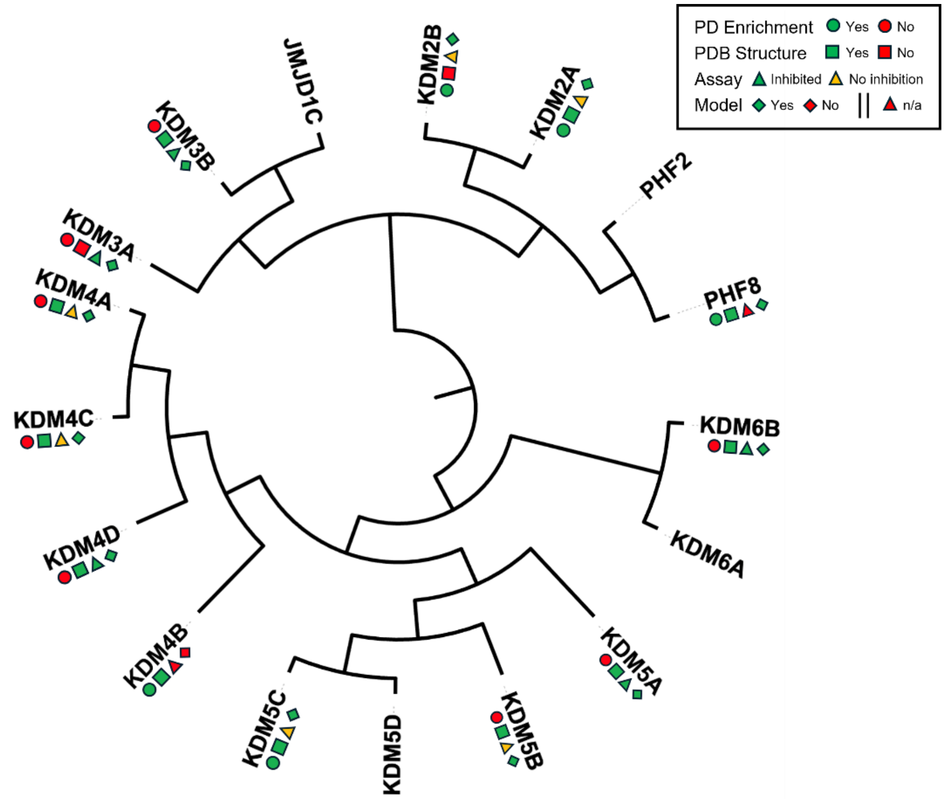


**Supplementary Fig. S3. Summary of observations and structural data reflected on the evolutionary relationships between JmjC domains.** The pictograms summarize the observed enrichment in one or more pull-down MS experiments (Fig. 2); the availability of X-ray crystal structures in the Protein Data Bank; whether inhibition was detected in one or more activity-based assays (50% inhibition at least <250 μM, Fig. S3); and if a Boltz2 model was generated (templated by an available X-ray crystal structure, Table S2). The tree represents the primary-sequence phylogenetic relationship of JmjC domain primary sequences calculated in ClustalW and plotted with the Interactive Tree of Life (ITOL) software[1,2].

**Table S4.** Activity-based demethylase assays from Reaction Biology Corporation (PA).

| Enzyme | Assay format | Substrate peptide | Substrate conc (nM) | αKG conc (μM) | PDCA $IC_{50}$ (μM) | GSK-J1 $IC_{50}$ (μM) |
|---|---|---|---|---|---|---|
| **KDM4A** | HTRF | H3 (21-44) K36Me3-Btn | 20 | 6 | 1.6 | – |
| **KDM4C** | HTRF | H3 (21-44) K36Me3-Btn | 50 | 5 | 0.18 | – |
| **KDM4D** | AlphaLisa | H3 (1-21) K9Me3-Btn | 25 | 5 | 0.1–0.2 | |
| **KDM5A** | HTRF | H3 (1-21) K4Me3-Btn | 25 | 12 | 1.1 | – |
| **KDM5B** | HTRF | H3 (1-21) K4Me3-Btn | 30 | 20 | 0.3 | – |
| **KDM5C** | HTRF | H3 (1-21) K4Me3-Btn | 10 | 15 | 1.1 | – |
| **KDM6B** | AlphaLisa | H3 (21-44) K27Me3-Btn | 60 | 10 | – | 0.003 |

CA = 2,4-pyridinedicarboxylic acid CAS 207671-42-9; GSK-J1 = inhibitor CAS 1373422-53-7; Btn=biotin.

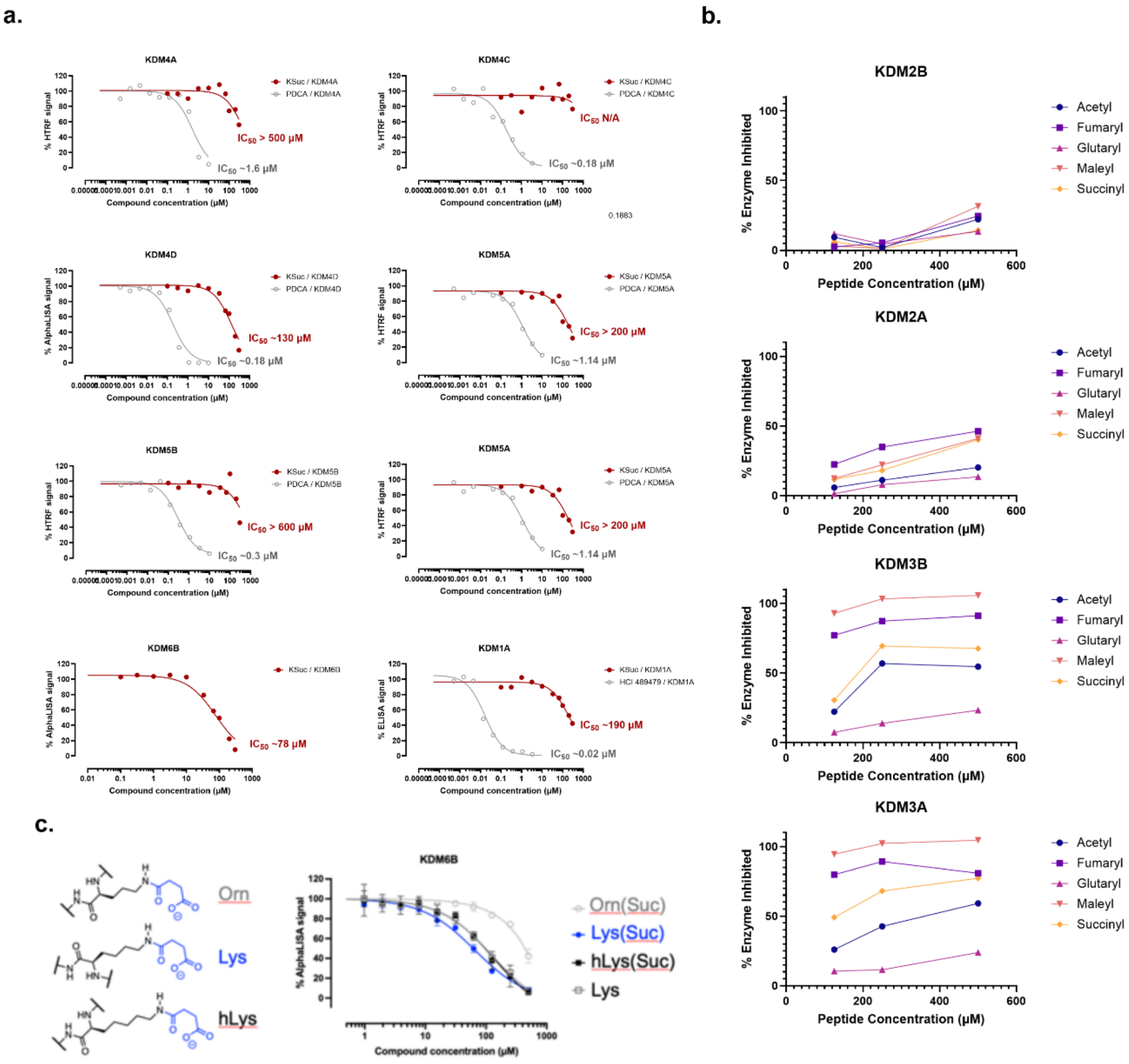


**Supplementary Fig. S4: Histone demethylase assays**. (a) Commercially available panel of histone demethylase assays tested in dose response with the peptide AARK(su)SA. The AlphaLISA and HTRF assays were done at Reaction Biology Corporation (Pennsylvania). (b) The LANCE histone demethylase assays were done at Eurofins (France).(c) Inhibition of KDM6B by unmodified lysine (negative control), homolysine, and ornithine.

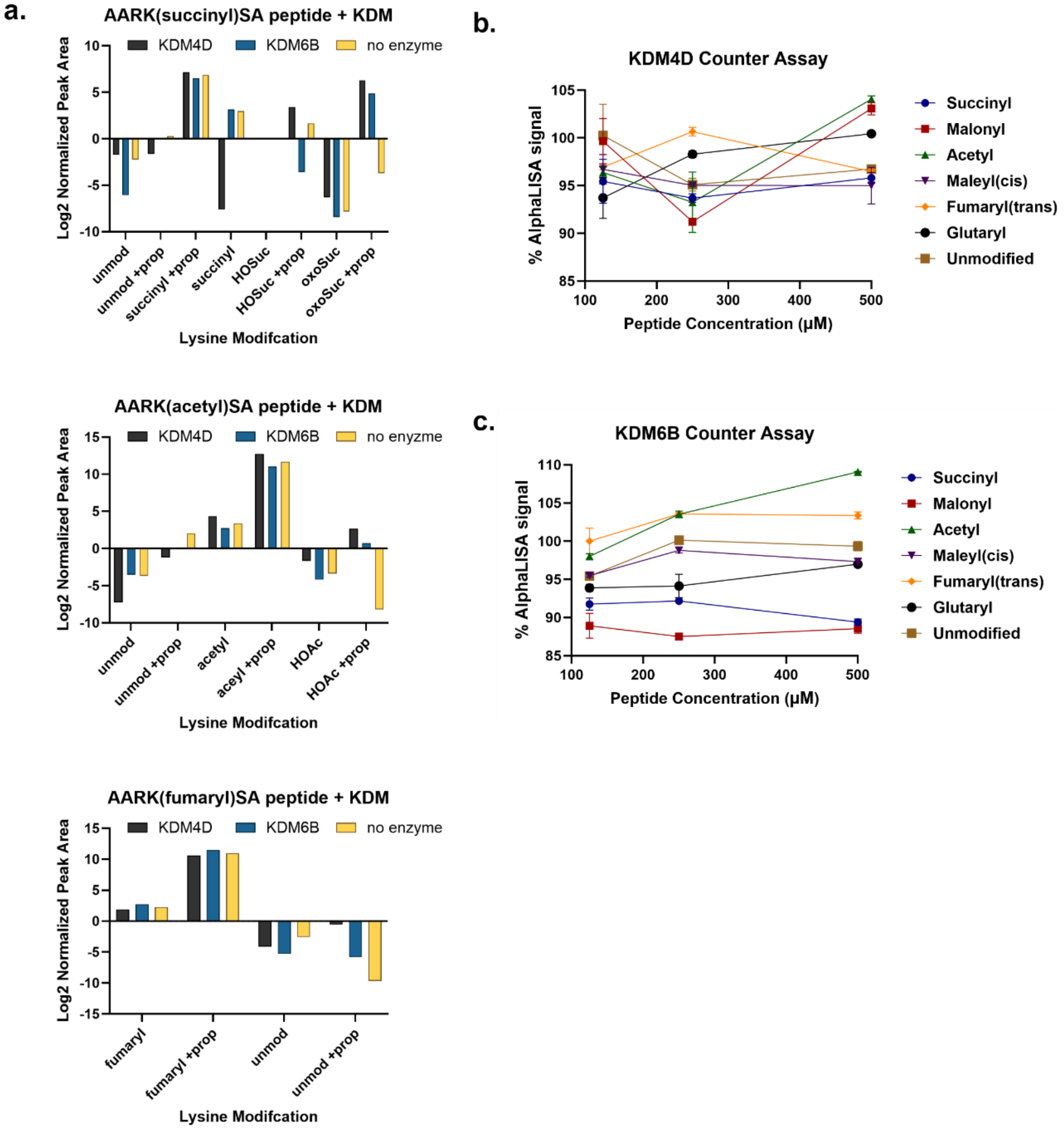


**Supplementary Fig. S5: Additional controls to validate the results of AlphaLISA assays.** (a) LC-mass spectrometry analysis of synthetic histone peptides used as inhibitor in AlphaLISA assay. Reaction products were analyzed to determine prevalence of de-acylation of peptides following enzyme incubation, or oxidation and hydrolysis of succinyl and acetyl groups. (b-c) AlphaLISA counter assay. Acylated synthetic peptides were incubated with synthetic di-methylated histone H3 peptides (reaction products), and then the AlphaLISA readout was performed to validate that these assays are free from any interference or fluorescence quenching by the Ksu and analog acyl-lysine peptides.

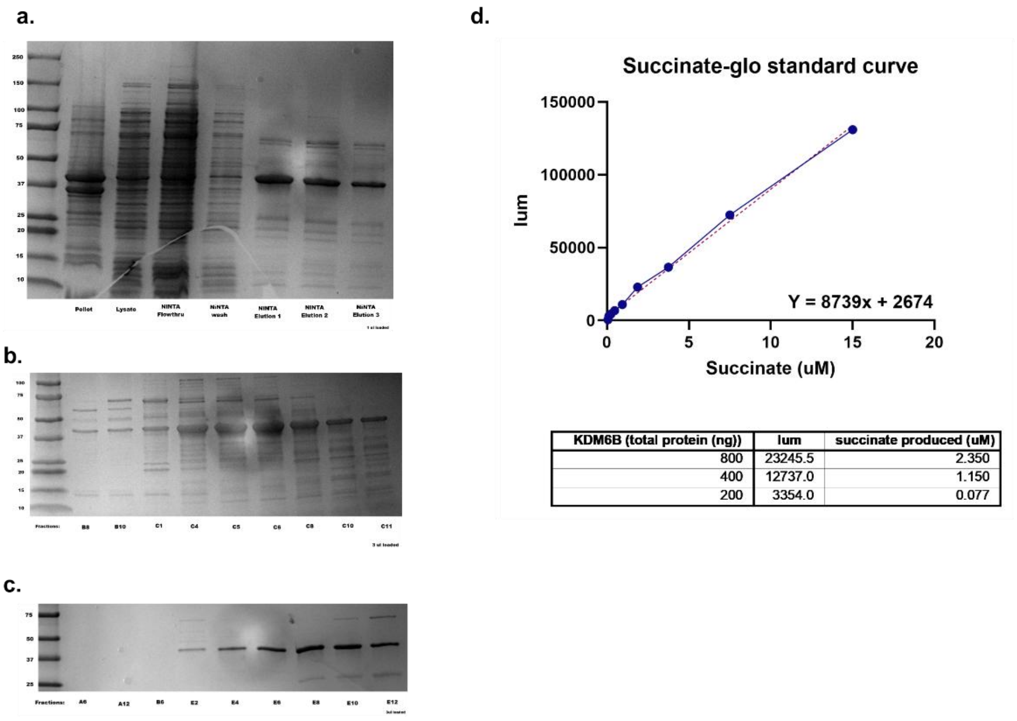


| KDM6B (total protein (ng)) | lum | succinate produced (uM) |
|---|---|---|
| 800 | 23245.5 | 2.350 |
| 400 | 12737.0 | 1.150 |
| 200 | 3354.0 | 0.077 |

**Supplementary Fig. S6: Purification and validation of recombinant KDM6B.** (a) SDS-PAGE gel of bacterial pellet, crude lysate, and fractions from NiNTA purification. (b) SDS-PAGE gel of fractions obtained from size exclusion chromatography. (c) SDS-PAGE gel of fractions obtained from ion exchange chromatography. (d) Results from succinate-glo assay. A dilution of a succinate standard was used to create a standard curve, and succinate production from a demethylase reaction performed with the purified KDM6B was calculated.

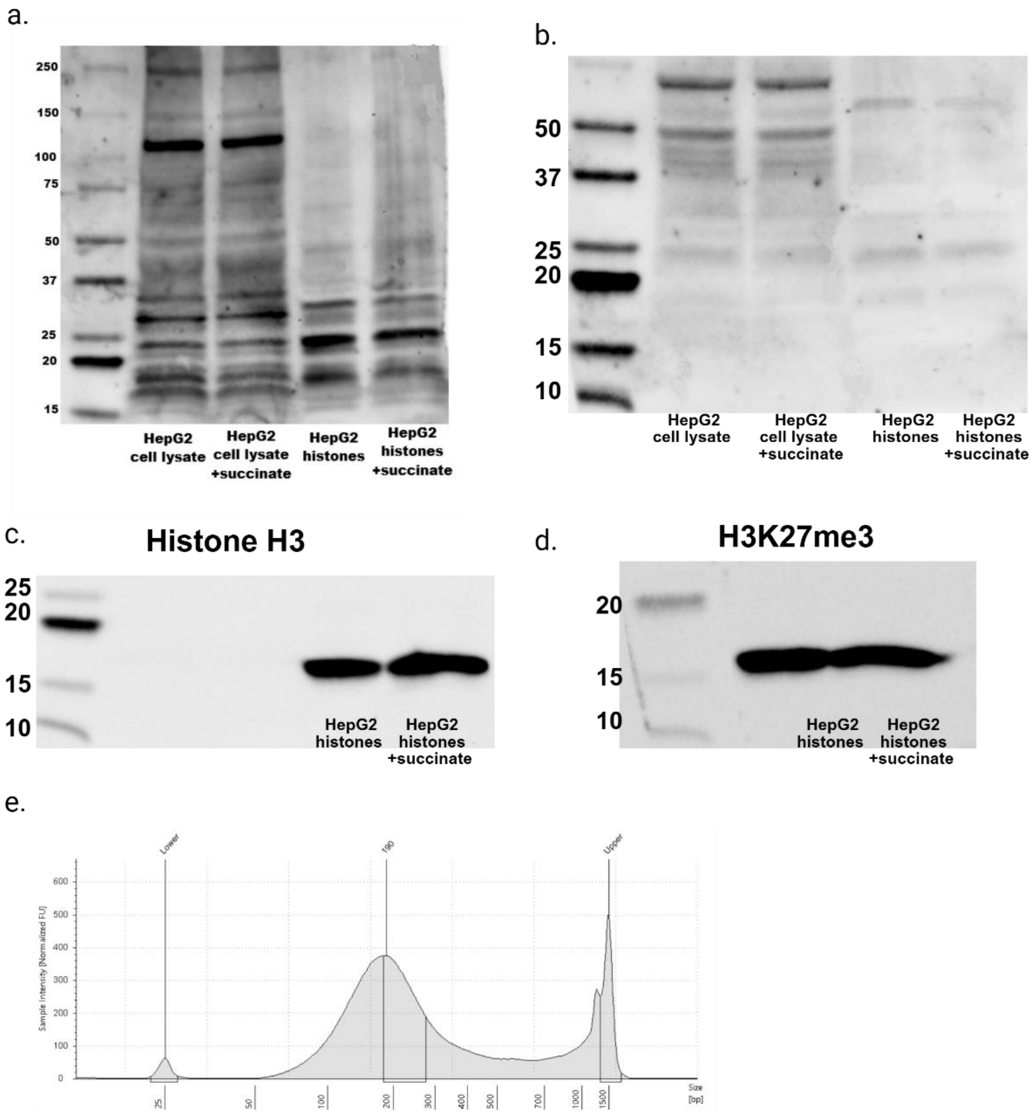


**Supplementary Fig. S7: Validation assays for ChIP-MS and CUT&Tag.** (a) Western blot validation of Succinylated Lysine Polyclonal Antibody (Invitrogen, PA5-120740) (CUT&Tag). (b) Western blot validation of PTMScan Succinyl-Lysine Motif [Succ-K] RmAb (Cell Signaling Technology) (ChIP-MS). (c) Western blot for histone H3 as part of validation scheme for Ksu antibodies Histone H3 (96C10) Mouse Monoclonal Antibody #3638, Cell Signaling Technology) (d) Western blot validation of Tri-Methyl-Histone H3 (Lys27) (C36B11) Rabbit Monoclonal Antibody #9733 (ChIP-MS assay). Full Western blot protocol is described in methods section.(e) Representative TapeStation (Agilent) trace of DNA fragment size following 16 cycles (30 sec on, 30 sec off) of sonication with Diagenode Pico sonicator.

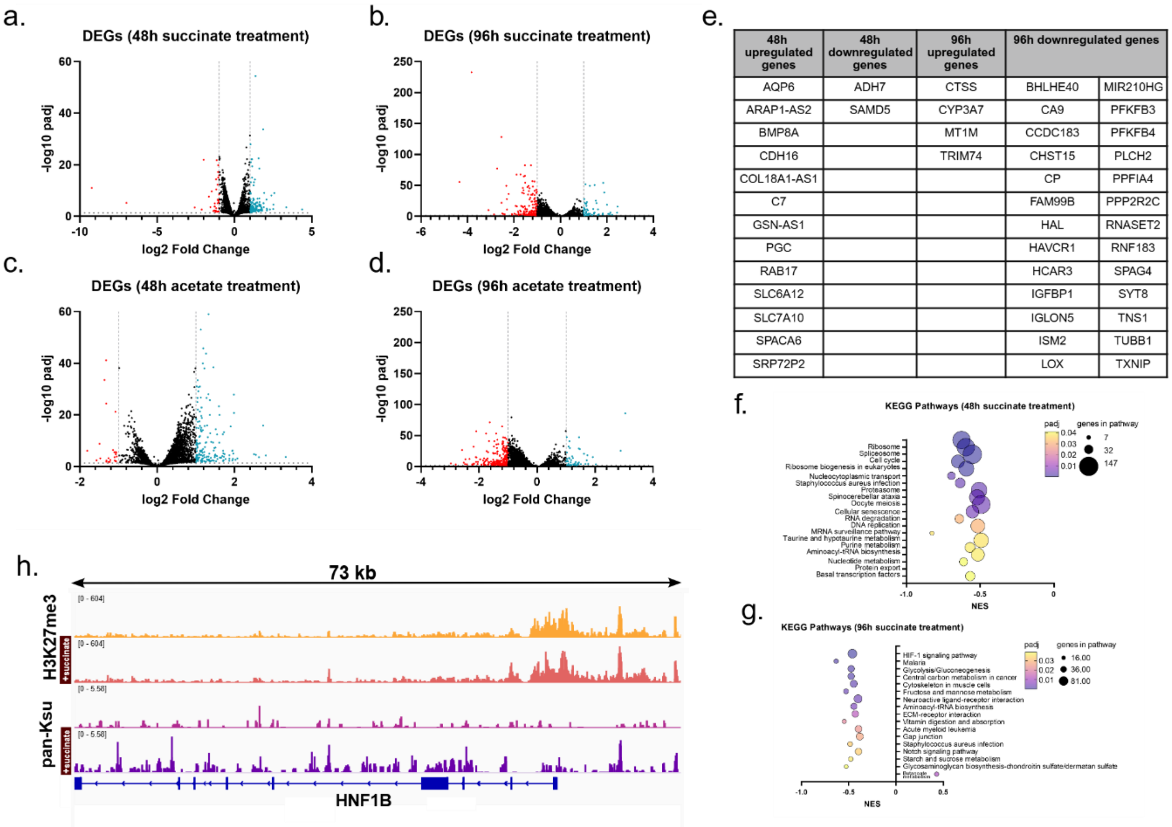


| 48h upregulated genes | 48h downregulated genes | 96h upregulated genes | 96h downregulated genes | |
|---|---|---|---|---|
| AQP6 | ADH7 | CTSS | BHLHE40 | MIR210HG |
| ARAP1-AS2 | SAMD5 | CYP3A7 | CA9 | PFKFB3 |
| BMP8A | | MT1M | CCDC183 | PFKFB4 |
| CDH16 | | TRIM74 | CHST15 | PLCH2 |
| COL18A1-AS1 | | | CP | PPFIA4 |
| C7 | | | FAM99B | PPP2R2C |
| GSN-AS1 | | | HAL | RNASET2 |
| PGC | | | HAVCR1 | RNF183 |
| RAB17 | | | HCAR3 | SPAG4 |
| SLC6A12 | | | IGFBP1 | SYT8 |
| SLC7A10 | | | IGLON5 | TNS1 |
| SPACA6 | | | ISM2 | TUBB1 |
| SRP72P2 | | | LOX | TXNIP |

**Supplementary Fig. S8: Additional transcriptomic analysis/CUT&Tag data.** (a) Volcano plot of DEGs following 48h succinate treatment, standard Deseq2 normalization. (b) Volcano plot of DEGs following 96h succinate treatment, standard Deseq2 normalization. (c) Volcano plot of DEGs following 48h acetate treatment, standard Deseq2 normalization. (d) Volcano plot of DEGs following 96h acetate treatment, standard Deseq2 normalization. (e) Table of significantly regulated genes shared between succinate and acetate treatment conditions. (f-g) KEGG pathway enrichment analysis of DEGs following succinate treatment. (FC threshold > 1 (abs), FDR < 0.05)

**Input for Boltz2 model**

Input 1: JmjC Domain (aa1141-1643, Selected as **Protein**)

VRASRNAKVKGKFRESYLSPAQSVKPKINTEEKLPREKLNPPTPSIYLESKRDAFSPV
LLQFCTDPRNPITVIRGLAGSLRLNLGLFSTKTLVEASGEHTVEVRTQVQQPSDENWDLTGTRQIWPCESSRSH
TTIAKYAQYQASSFQESLQEEKESEDEESEEPDSTTGTPPSSAPDPKNHHIIKFGTNIDLSDAKRWKPQLQELL
KLPAFMRVTSTGNMLSHVGHTILGMNTVQLYMKVPGSRTPGHQENNNFCSVNINIGPGDCEWFAVHEHYWETIS
AFCDRHGVDYLTGSWWPILDDLYASNIPVYRFVQRPGDLVWINAGTVHWVQATGWCNNIAWNVGPLTAYQYQLA
LERYEWNEVKNVKSIVPMIHVSWNVARTVKISDPDLFKMIKFCLLQSMKHCQVQRESLVRAGKKIAYQGRVKDE
PAYYCNECDVEVFNILFVTSENGSRNTYLVHCEGCARRRSAGLQGVVVLEQYRTEELAQAYDAFTLAPASTSR

Input 2: Fe2+ atom (Selected as **Ligand**)

CCD Code: FE2

Input 3: Succinyl Lysine Peptide (AARK(Suc)AA sequence, Selected as **Ligand**)

SMILES:
O=C([C@H](CCCNC(N)=[NH2+])NC([C@H](C)NC([C@H](C)NC(C)=O)=O)=O)N[C@@H](CCCC
NC(CCC([O-])=O)=O)C(N[C@H](C(N[C@H](C(N)=O)C)=O)C)=O

Input 4: Water (Selected as **Ligand**)

CCD Code: HOH

Input 5: Water (Selected as **Ligand**)

CCD Code: HOH
